# Tip rates, phylogenies, and diversification: what are we estimating, and how good are the estimates?

**DOI:** 10.1101/369124

**Authors:** Pascal O. Title, Daniel L. Rabosky

## Abstract

1. Species-specific diversification rates, or “tip rates”, can be computed quickly from phylogenies and are widely used to study diversification rate variation in relation to geography, ecology, and phenotypes. These tip rates provide a number of theoretical and practical advantages, such as the relaxation of assumptions of rate homogeneity in trait-dependent diversification studies. However, there is substantial confusion in the literature regarding whether these metrics estimate speciation or net diversification rates. Additionally, no study has yet compared the relative performance and accuracy of tip rate metrics across simulated diversification scenarios.
2. We compared the statistical performance of three model-free rate metrics (inverse terminal branch lengths; node density metric; DR statistic) and a model-based approach (BAMM). We applied each method to a large set of simulated phylogenies that had been generated under different diversification processes; scenarios included multi-regime time-constant and diversity-dependent trees, as well as trees where the rate of speciation evolves under a diffusion process. We summarized performance in relation to the type of rate variation, the magnitude of rate heterogeneity and rate regime size. We also compared the ability of the metrics to estimate both speciation and net diversification rates.
3. We show decisively that model-free tip rate metrics provide a better estimate of the rate of speciation than of net diversification. Error in net diversification rate estimates increases as a function of the relative extinction rate. In contrast, error in speciation rate estimates is low and relatively insensitive to extinction. Overall, and in particular when relative extinction was high, BAMM inferred the most accurate tip rates and exhibited lower error than non-model-based approaches. DR was highly correlated with true speciation rates but exhibited high error variance, and was the best metric for very small rate regimes.
4. We found that, of the metrics tested, DR and BAMM are the most useful metrics for studying speciation rate dynamics and trait-dependent diversification. Although BAMM was more accurate than DR overall, the two approaches have complementary strengths. Because tip rate metrics are more reliable estimators of speciation rate, we recommend that empirical studies using these metrics exercise caution when drawing biological interpretations in any situation where the distinction between speciation and net diversification is important.

## Introduction

Rates of speciation and extinction vary through time and among lineages (Nee, Mooers & Harvey 1992; Sanderson & Donoghue 1996; Etienne & Haegeman 2012; Jetz et al. 2012; Moen & Morlon 2014; Alfaro et al. 2018), contributing to dramatic heterogeneity in species richness across the tree of life (Alfaro et al. 2009; Jetz et al. 2012; Barker et al. 2013). By characterizing variation in rates of speciation and extinction, we can better understand the dynamics of biological diversity through time, across geographic and environmental gradients (Zink, Klicka & Barber 2004; Ricklefs 2006; Mittelbach et al. 2007; Silvestro, Schnitzler & Zizka 2011; Rabosky, Title & Huang 2015), and in relation to traits and key innovations (FitzJohn, Maddison & Otto 2009; Near et al. 2012; Beaulieu & O’Meara 2016). Consequently, there has been great interest in statistical methods for inferring rates of speciation and extinction from molecular phylogenies.

Although rates of diversification have traditionally been quantified for clades, there has been a growing interest in estimating species-specific rates of diversification, which we refer to here as “tip rates”. Tip rates are increasingly used to describe patterns of geographic and trait-associated variation in diversification (Freckleton, Phillimore & Pagel 2008; Jetz et al. 2012; Kennedy et al. 2016; Harvey & Rabosky 2017; Quintero & Jetz 2018; Rabosky et al. 2018). It may seem strange to view evolutionary rates as a property of individual lineages, but such rates emerge naturally from the birth-death model we typically use to conceptualize the diversification process (Nee, Mooers & Harvey 1992; Nee, May & Harvey 1994). Under the birth-death process, individuals (species) are characterized by per-lineage rates of species origination (speciation, λ) and extinction (μ). For the purposes of inference, these rates are typically assumed to be constant among contemporaneous members of a focal clade. However, tip rates can be viewed as our best estimate of the present-day rate of speciation or extinction for an individual lineage, conditional on past (usually recent) evolutionary history. As such, they provide information about the expected amount of time that will elapse before a lineage splits or becomes extinct.

A number of approaches have been used to estimate tip rates, including both model-based and non-model-based approaches (i.e., models that are parameterized with speciation and extinction rates, vs metrics that simply rely on branch lengths and splitting events). These approaches vary in terms of how much information they derive from a focal species (i.e., a terminal branch) relative to the amount of information they incorporate from other regions of the phylogeny. On one end of the spectrum, tree-wide estimates (i.e., one rate for the entire phylogeny) of speciation and extinction rates under a constant-rate birth-death (CRBD) model provide tip rates that are maximally auto-correlated (identical) across species in the clade; such rates for any given species are not independent of rates for any other species in the group of interest. On the other end of the spectrum, terminal branch lengths can be used to derive a censored estimate of the rate of speciation that is minimally autocorrelated with rates for other species in the focal clade. Terminal branch lengths are largely unique to each species (rates might be identical only for sister taxa), but provide a noisy measure of speciation, due to the stochasticity inherent in the diversification process (Nee, May & Harvey 1994), and they have been employed as a summary statistic in assessing model adequacy (Bromham, Hua & Cardillo 2016; Gomes, Sorenson & Cardoso 2016). In contrast to single (terminal) branch estimates, tree-wide estimates should be less susceptible to stochastic noise, because they incorporate information from the entirety of the tree (e.g., multiple branches are used in the estimates). Of course, the tree-wide estimate necessarily assumes that all tips share a common underlying diversification process. Other tip rate metrics fall somewhere between these two extremes, incorporating some tree-wide information but relaxing the assumption of homogeneous rates across all lineages (node density metric: Freckleton, Phillimore & Pagel 2008; DR: Jetz et al. 2012). The estimation of tip-specific rates thus entails a tradeoff between the precision of individual estimates and the stochastic error associated with those estimates.

BAMM (Bayesian Analysis of Macroevolutionary Mixtures, Rabosky 2014) is a model-based approach that can accommodate heterogeneity in the rate of diversification through time and among lineages. BAMM simulates a posterior distribution of macroevolutionary rate shift configurations given a phylogeny of interest; marginal rates of speciation and extinction for individual taxa can then be extracted from this distribution. In this framework, the correlation in rates between any pair of species is a function of the posterior probability that they share a common macroevolutionary rate regime (Rabosky et al. 2014). If the tree-wide posterior probability of rate variation is low, the marginal rates estimates for individual species will be similar across the entire tree, as under a CRBD model. Likewise, any pair of taxa that are consistently assigned to the same macroevolutionary rate regime will necessarily have identical tip rates.

Tip rates are best suited to a host of questions and hypotheses where the diversification dynamics over the evolutionary history of a group are either less relevant, or no more relevant, than the rates of diversification closer to the present day. For example, many hypotheses involving trait-dependent diversification implicitly assume a time-homogeneous, or constant through time, effect of the trait on diversification rate (Coyne & Orr 2004; Kay et al. 2006; Jablonski 2008; FitzJohn 2010; Claramunt et al. 2011). Harvey & Rabosky (2017) found that the use of tip rates for assessing correlations between continuous traits and diversification has good performance across a range of diversification scenarios. Furthermore, hypotheses pertaining to non-historical geographic patterns of diversity are also better addressed with recent rates of diversification. For example, many hypotheses for the latitudinal diversity gradient propose time-homogeneous effects of particular environmental factors (temperature, energy, geographic area) on rates of diversification (Mittelbach et al. 2007; Kennedy et al. 2014; Rabosky, Title & Huang 2015; Schluter 2016; Rabosky et al. 2018). Put simply, if such time-homogeneous processes have shaped the latitudinal diversity gradient (e.g., correlation between speciation and temperature: Rohde 1992), then the effect should be manifest in the distribution of present-day evolutionary rates.

At present, there is substantial confusion in the literature over what quantity various tip rate metrics actually measure. The DR statistic (Jetz et al. 2012) was originally described as a measure of the “species-level lineage diversification rate”. While supplemental analyses and subsequent work suggested that DR was a better measure of speciation rate than net diversification (Jetz et al. 2012, Belmaker & Jetz 2015, Quintero & Jetz 2018), many studies have nonetheless continued to describe DR as an estimate of the lineage-level net diversification rate (Marin & Hedges 2016; Oliveira et al. 2016; Cai et al. 2017; and many others). The node density metric of Freckleton, Phillimore & Pagel (2008) has also been described as a measure of net diversification. Whether these metrics more accurately measure speciation or net diversification is critically important for interpreting biodiversity patterns (e.g., two regions might differ dramatically in speciation rate, but net diversification rates in each might nonetheless be zero). An objective of our study is thus to compare the ability of DR, node density, and other metrics to estimate speciation and net diversification rates.

Despite the potential utility of tip rates in geographic and trait-based analyses of speciation rate heterogeneity (Jetz et al. 2012; Belmaker & Jetz 2015; Oliveira et al. 2016; Quintero & Jetz 2018), there has yet been no comprehensive comparative assessment of the accuracy and precision of the estimates, save for supplemental analyses in Jetz et al. (2012) and Quintero & Jetz (2018). BAMM has low power to infer small rate regimes (Rabosky, Mitchell & Chang 2017; Meyer & Wiens 2017), leading to the possibility that other approaches might perform better for smaller phylogenies or when the variation in rates among clades is subtle. However, DR and related methods will always identify variation in tip rates, even when none exists, provided there is stochastic variation in branch lengths. A goal of this study is therefore to evaluate the trade-off between the stochastic noise inherent in non-model-based approaches, and the conservative but less noisy estimates from model-based metrics. We compare the performance of these metrics across a range of simulation scenarios, which include both discrete and continuous variation in rates.

## Methods

### Tip rate metrics

We assessed the accuracy of four tip rate metrics in this study at quantifying rates of speciation. As we demonstrate below (see also Supplementary Figure 5 in Jetz et al. 2012; extended Figure 5 in Quintero & Jetz 2018; Belmaker & Jetz 2015), these metrics are estimators of speciation rate and not net diversification rate, and we refer to them as such throughout. The first metric is the inverse of the equal splits measure (Redding and Mooers 2006), also called the *DR statistic* (Jetz et al. 2012), *DivRate* (Belmaker & Jetz 2015; Oliveira et al. 2016), *ES* (Harvey & Rabosky 2017) or *tip DR* (Quintero & Jetz 2018), which we denote in this study as β_DR_. This species-specific measure incorporates the number of splitting events and the internode distances along the root-to-tip path of a phylogeny, while giving greater weight to branches closer to the present (Redding & Mooers 2006; Jetz et al. 2012). β_DR_ is computed as:

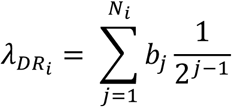

where 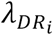 is the tip rate for species *i, N*_*i*_ is the number of branches between species *i* and the root, *b*_*j*_ is the length of branch *j*, starting at the terminal branch (*j* = 1) and ending with the root. Jetz et al. (2012) demonstrated that, for trees deriving from a Yule process, and with mild extinction, the mean λ_DR_ across tips converges on the true speciation rate.

We also considered a simpler metric, node density (Freckleton, Phillimore & Pagel 2008; denoted by λ_ND_). This is simply the number of splitting events along the path between the root and tip of a phylogeny, divided by the age of the phylogeny. While λ_DR_ down-weights the contribution of branch lengths that are closer to the root, λ_ND_ equally weights the contributions of all branches along a particular root-to-tip path, regardless of where they occur in time. Under a pure-birth model (μ = 0), both λ_DR_ and λ_ND_ should yield unbiased estimates of the rate of speciation.

The third measure we considered is the inverse of the terminal branch lengths (λ_TB_). Rapid speciation rates near the present should be associated with proportionately shorter terminal branches; smaller values of λ_TB_ should thus characterize species with faster rates of speciation. This measure has recently been used as a summary statistic to assess model adequacy in trait-dependent diversification studies (Bromham, Hua & Cardillo 2016; Gomes, Sorenson & Cardoso 2016; Harvey & Rabosky 2017). Following Steel & Mooers (2010), we note that the terminal branch lengths can be used to derive an estimate of the speciation rate; this follows from the fact that interior and terminal branches have the same expected value under the Yule process (Steel & Mooers 2010). The corresponding estimator for the *i*’th tip, λ_TB_ is approximately 1 / 2*b* where *b* is the length of a given terminal branch (Steel & Mooers 2010). To our knowledge, λ_TB_ has not been used to explicitly estimate tip rates as we do here, but given its utility as a summary statistic and general theoretical properties (Steel & Mooers 2010), we see value in comparing the performance of this metric to others currently in use.

Finally, we considered a Bayesian, model-based approach to estimating tip rates. BAMM (Rabosky 2014) assumes that phylogenies are generated by set of discrete diversification regimes. Using MCMC, the program simulates a posterior distribution of rate shift regimes, from which marginal posterior rate distributions can be extracted for each tip in the phylogeny. Priors for BAMM analyses were set using default settings from the setBAMMpriors function from BAMMtools (Rabosky et al. 2014). The prior parameterizations specified by this function ensure that the prior density on relative rate changes across the tree is invariant to the scale of the tree (e.g., multiplying branch lengths by 106 will not change inferences about relative rates across the tree). We denote BAMM tip speciation rates (mean of the marginal posterior) as λ_BAMM_. As BAMM also estimates extinction rates for each regime, we also calculated tip-specific net diversification rate as λ_BAMM_ − μ_BAMM_, denoted as *r*_BAMM_.

### Tip rate metrics estimate speciation, not net diversification

As suggested previously (Belmaker & Jetz 2015; supplemental analyses in Jetz et al. 2012), DR and presumably other tip-based measurements, more accurately estimate the rate of speciation than the rate of net diversification. However, numerous studies continue to refer to DR as a measure of net diversification (Marin & Hedges 2016; Oliveira et al. 2016; Cai et al. 2017; Quintero & Jetz 2018; and many others). This is incorrect and it is straightforward to demonstrate that λ_TB,_ λ_ND_ and λ_DR_ are more reliable measures of speciation rates and not net diversification rates, at least when extinction is moderate to high.

To illustrate this property of the metrics, we applied all approaches to constant-rate birth-death phylogenies simulated across a range of extinction fractions (ε = μ /λ), including pure-birth trees (ε = 0) as well as trees exhibiting very high turnover (ε = 1). To evaluate accuracy of speciation estimates as a function of ε, we generated 1000 phylogenies with 100 tips each, where λ and ε were drawn from uniform distributions (λ: [0.05, 0.3]; ε: [0, 1]). Importantly, when λ is sampled uniformly with respect to ε, the distribution of *r* is not uniform: the mean, range and variance in *r* decrease dramatically as ε increases. To evaluate the accuracy of *r* as a function of ε, we thus generated a second set of trees by sampling *r* and ε from uniform distributions (*r*: [0.05, 0.3], ε [0, 1]). As a result, λ has constant mean and variance with respect to ε in the first set of simulations, and the same is true for *r* in the second set of simulations (Figure S1). All phylogeny simulations were conducted with the TreeSim package in R (Stadler 2011).

We compared tip rate metrics to true speciation rates λ_TRUE_ (with the first simulation set) and to true net diversification rates *r*_TRUE_ (with the second simulation set). We evaluated mean per-tip accuracy of the tip rate metrics with three measures of error:

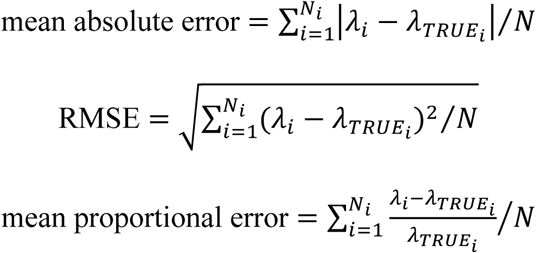

where λ_i_ is the estimated tip rate for species *i* out of N total species, λ_TRUE_ is the true tip rate. Mean absolute error and root mean square error capture the magnitude in error in tip rates, and mean proportional error quantifies the bias in tip rates, as a function of the true tip rates (Rabosky et al. 2014). In analyses below, all error summaries yield generally congruent results; results for mean absolute error are presented in the main text, and others in supplement.

### Assessment of tip rate metrics

We tested the performance of the metrics by compiling publicly-available datasets from a number of simulation-based studies (Table 1). By focusing on simulations from previously-published work, we thus ensured that the simulation process itself was effectively blinded to the objectives of the present study. We further note that our trial datasets included several studies that were critical of BAMM (Moore et al. 2016, Meyer & Wiens 2017). These simulated trees include rate heterogeneity in time and across lineages. Together, these phylogenies present a wide range of tree sizes and diversification rate shifts, providing an ideal comparative dataset for our purposes. To more easily distinguish between these tree types in the text, we refer to the BAMM-type, multi-regime time-constant phylogenies simply as “multi-regime”, and the multi-regime diversity-dependent phylogenies simply as “diversity-dependent”, even though discrete rate shifts are present in both types of trees. In addition to discrete-shift scenarios (e.g., BAMM-type process), we simulated phylogenies under an “evolving rates” model of diversification (Rabosky 2010; as corrected in Beaulieu & O’Meara 2015) to explore performance of tip rate metrics when diversification rates change continuously and independently along branches, as might occur if diversification rates are correlated with an underlying continuous trait (FitzJohn 2010). In these simulations, we allowed the logarithm of λ to evolve across the tree under a Brownian motion process, while holding ε constant. The magnitude of rate heterogeneity among branches is controlled by the diffusion parameter σ, where greater values lead to greater heterogeneity in speciation rates. Although published phylogenies with rate data were unavailable for this simulation scenario, we used simulation code and parameters taken directly from Beaulieu & O’Meara (2015) to generate trees with similar statistical properties to those in their study. Simulations were performed with the following parameters: λ = 0.078, 0.103, 0.145, 0.249 and ε = 0.0, 0.25, 0.50, 0.75. We simulated 100 phylogenies for each (λ, ε) pair, and for three values of σ (σ = 0.03, 0.06, 0.12). We evaluated tip rate accuracy by comparing estimated to true tip rates, using the absolute and proportional error metrics described above. We also examined the correlation between true and estimated tip rates, combining tip rates from all phylogenies generated under the same class of diversification process, and visualizing these data as density scatterplots, generated with the LSD package in R (Schwalb et al. 2018), where colors indicate the density of points.

**Table 1.** Summary of simulated phylogenies used in this study.

| simulation model | number<br>of trees | tree size | regime<br>number | source |
| --- | --- | --- | --- | --- |
| single-regime, constant-rate birth-death | 100 | 100 | 1 | Mitchell & Rabosky 2016 |
| single- and multi-regime, constant-rate birth-death | 100 | 51-148 | 1-6 | Moore et al. 2016 |
| single- and multi-regime, constant-rate birth-death | 400 | 10-4296 | 1-67 | Rabosky, Mitchell & Chang 2017 |
| multi-regime, constant-rate birth-death | 20 | 939-3708 | 11 | Meyer & Wiens 2017 |
| single- and multi-regime, constant-rate birth-death | 188 | 4-3955 | 1-73 | Mitchell, Etienne & Rabosky 2018 |
| single-regime, constant-rate birth-death, lambda uniform | 1000 | 100 | 1 | this study |
| single-regime, constant-rate birth-death, net diversification uniform | 1000 | 100 | 1 | this study |
| pure birth root regime, 1-4 discrete shifts to diversity-dependent regimes | 1200 | 54-882 | 1-5 | Rabosky 2014; Mitchell & Rabosky 2016 |
| speciation rate evolves via diffusion process | 1200 | 25-1208 | 1 | Rabosky 2010; Beaulieu & O'Meara 2015; Rabosky 2016; this study |

Size of diversification rate regimes might be an important factor in a tip rate metric’s ability to accurately estimate rates. For example, BAMM’s statistical power in detecting a shift to a new rate regime is a function of the number of taxa in that rate regime, and tip rates for taxa from small regimes will more likely be parameterized according to the larger parent regime or the tree-wide average rate (Rabosky, Mitchell & Chang 2017); this is the expected behavior when BAMM fails to identify a rate shift. However, non-model-based approaches such as those examined in this study might be more accurate for small regimes. To explore how rate regime size influences the accuracy of tip rate metrics, we calculated the mean tip rate for each true rate regime from all multi-regime phylogenies (simulation datasets from Moore et al. 2016; Rabosky, Mitchell & Chang 2017; Meyer & Wiens 2017; Mitchell, Etienne & Rabosky 2018). We then calculated the Pearson correlation coefficient and the slope of a linear model between true and estimated mean regime rates. We explored the performance of all metrics with respect to regime sample size, as in Rabosky, Mitchell & Chang (2017: Figure 13). For comparison, we repeated all performance summaries on tip rates estimated by applying a simple constant-rate birth-death (CRBD) process to each simulated phylogeny. This exercise is an important control, because it indicates how much error we would expect for each simulated phylogeny under the simplifying (incorrect) assumption that rates are constant among lineages and through time for each dataset.

## Results

### Speciation or net diversification?

As expected, the tip rate metrics examined in this study are more accurate estimators of the rate of speciation (λ) and not the net rate of species diversification (*r*). Mean absolute error increased exponentially with respect to the extinction fraction ε (Figure 1). However, mean absolute error in speciation rate was largely invariant with respect to ε (0.95 quantile of *r*-based and λ-based mean absolute error for λ_DR_: 2.28 and 0.17, respectively). Nearly identical patterns were found with RMSE (Figure S2). Note that *r* and λ for these simulations were drawn from identical uniform distributions, and absolute error in the rates is thus comparable. Proportional error generally exhibited the same pattern, and in terms of λ versus *r*, differences in speciation-based error varied across ε (Figure S3). There was a weak but significant trend towards progressively greater underestimation of speciation rates with increasing values of relative extinction (linear model slopes: −0.08, −0.014, −0.011 for λ_ND_, λ_DR_ and λ_BAMM_, respectively). Overall, error was highest for λ_TB_ by an order of magnitude (Figure S4), and decreased progressively with λ_ND_ and λ_DR_, with the lowest overall error in λ_BAMM_. BAMM estimates of net diversification rate were relatively accurate, except at the highest values of ε (Figures 1, S2, S3).

**Figure 1.**
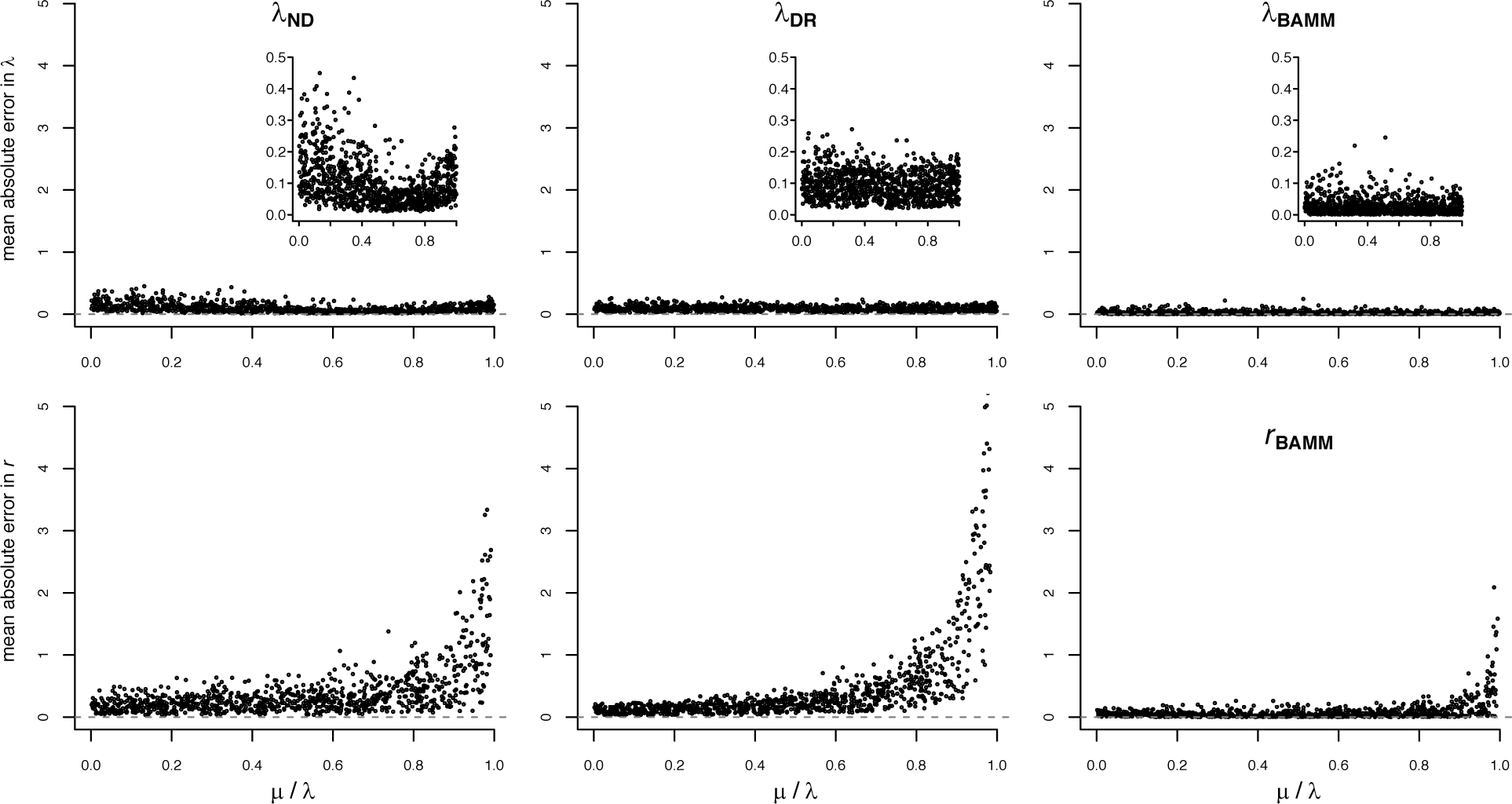
Mean absolute error in λ (top) and *r* (bottom) for three different tip rate metrics, across a range of relative extinction rates. For BAMM, the estimated speciation and net diversification rates are presented in the top and bottom panels, respectively. Absolute error of zero implies perfect accuracy. Inset plots show error in λ with truncated y-axis scale to facilitate comparison among metrics. All tip rate metrics track λ more accurately than they track *r*. See Figure S4 for λ_TB_, which performed much worse than the other metrics.

### Tip rate accuracy across rate-variable phylogenies

Tip rates estimated with BAMM were consistently more accurate than those obtained using the other methods across all diversification scenarios considered, including multi-regime, diversity-dependent and evolving rates trees (Figure 2). λ_DR_ was the second-most accurate metric, although its relationship with true rates was substantially weaker than λ_BAMM_. λ_ND_ and λ_TB_ were correlated with true rates but performed relatively poorly overall. However, λ_TB_ performed better than λ_ND_, and just as well as λ_DR_ at estimating speciation rates for diversity-dependent trees (Figure 2, S5). All metrics performed best for multi-regime trees, followed by evolving rates and diversity-dependent trees, respectively. For diversity-dependent trees, λ_ND_ rates are effectively uncorrelated with the true rates (Figure 2). Additionally, the performance of the different tip rate metrics for multi-regime phylogenies is not sensitive to the source of the simulated phylogenies (Figure S6). We found that BAMM substantially outperformed all other metrics on datasets from studies that independently assessed BAMM’s performance (Figure S6: Moore et al. 2016; Meyer & Wiens 2017). Tip rates were also generally but more weakly correlated with true net diversification rates, with the exception of λ_ND_, which was uncorrelated with true rates for diversity-dependent trees, presumably because this metric equally weights the full depth of the tree (Figure S7).

**Figure 2.**
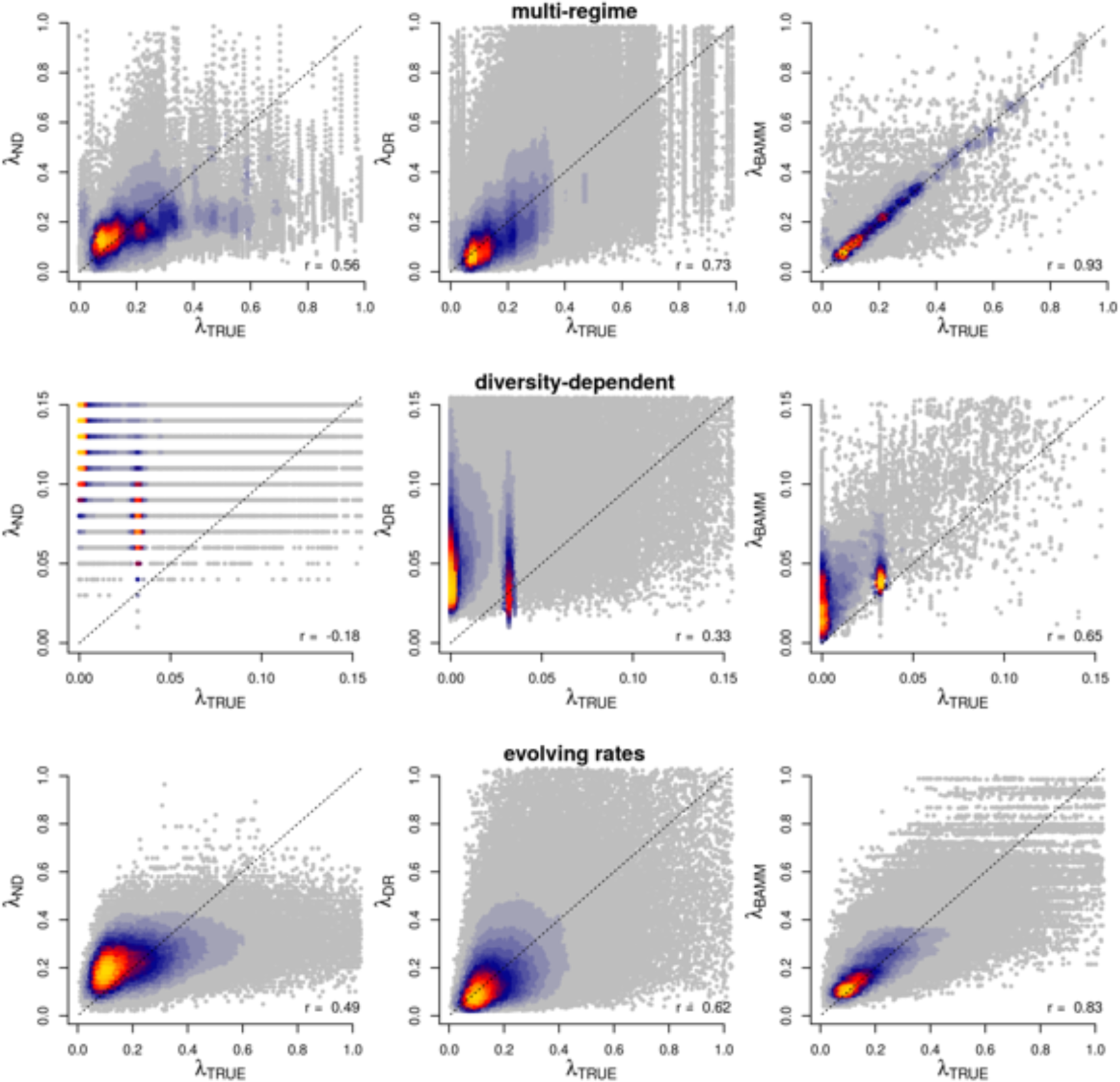
True tip rates (λ_TRUE_) in relation to estimated tip rates. Tip rates were compared separately for different major categories of phylogeny simulations (rows) and are plotted separately by inference method (columns). Plotting region is restricted to the 99th percentile of true rates, but Spearman correlations between true and estimated rates (lower right of each figure panel) are based on the full range of the data. Colors indicate the density of points in the scatter plots. The horizontal gaps in λ_ND_ for diversity-dependent trees are an artefact of all trees having the same crown age. λ_BAMM_ exhibited the strongest correlation with true rates for all simulation categories.

In terms of mean per-tip error, λ_BAMM_ consistently outperformed the other metrics for multi-regime, diversity-dependent and evolving rates trees (Figures 3, S8). Error in λ_BAMM_ increased as a function of rate heterogeneity for evolving rate phylogenies, but was largely independent of the magnitude of rate heterogeneity for the other scenarios. λ_DR_ generally exhibited greater error than λ_BAMM_, and this error increased as a function of the level of heterogeneity for both the evolving rates and multi-regime trees. Error in λ_DR_ was generally invariant to the number of rate regimes for the diversity-dependent scenarios. However, λ_DR_ tended to have greater error than tip estimates from a simple model that assumes no variation in rates through time or among lineages (λ_CRBD_; all tips assigned the tree-wide CRBD rate). λ_ND_ performed somewhat similarly to λ_DR_ for constant-rate and evolving rates trees, but worse for diversity-dependent trees. Error in λ_TB_ increased with increasing rate heterogeneity for constant-rate and evolving rates trees, but was relatively unaffected by rate heterogeneity in diversity-dependent trees (Figure S9). However, error for this metric was far greater than for all other tip metrics.

**Figure 3.**
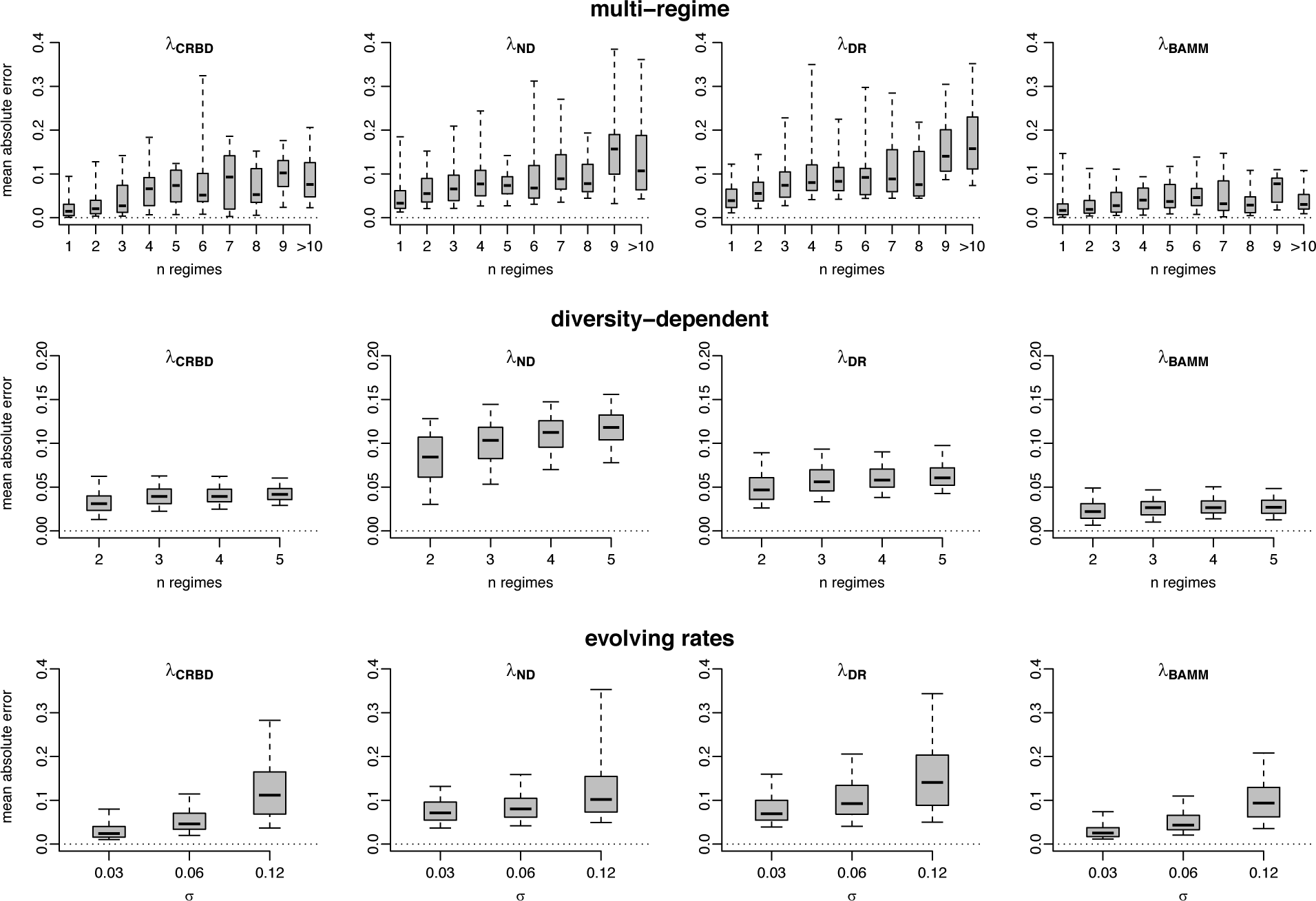
Mean per-tip absolute error in speciation rates as a function of the magnitude of rate heterogeneity in each simulated phylogeny. Results are presented separately for different categories of rate variation (Table 1); left column shows estimates from a constant-rate birth-death model for reference. The boxes and whiskers represent the 0.25 – 0.75, and the 0.05 – 0.95 quantile ranges, respectively. In some cases, λ_ND_ and λ_DR_ had more error than a simple CRBD model with no variation in tip rates. λ_BAMM_ had the least amount of error across all amounts of rate heterogeneity. See Figure S9 for λ_TB_.

### Effects of regime size on performance

Both metrics of performance assessment – the Pearson correlation and OLS slope – generally increased with increasing regime size (Figure 4). This was found to be true for all tip rate metrics, although λ_TB_ and λ_ND_ never achieved high performance. λ_DR_ tended to perform better than other metrics when small rate regimes were included (e.g., 10 tips or fewer); however, the slope between estimated and true rates was greater than 1 across the majority of minimum regime sizes, indicating that λ_DR_ overestimates speciation rates (see also Figure S3). Similar patterns were observed for net diversification rates with λ_DR_, but the magnitude of the overestimation was greater than for speciation (Figure S10). λ_BAMM_, in contrast, approached a slope of 1 when estimating speciation rates and slightly underestimated net diversification rates (regimes with > 30 tips: OLS slope = 0.96 for λ, 0.87 for *r*).

**Figure 4.**
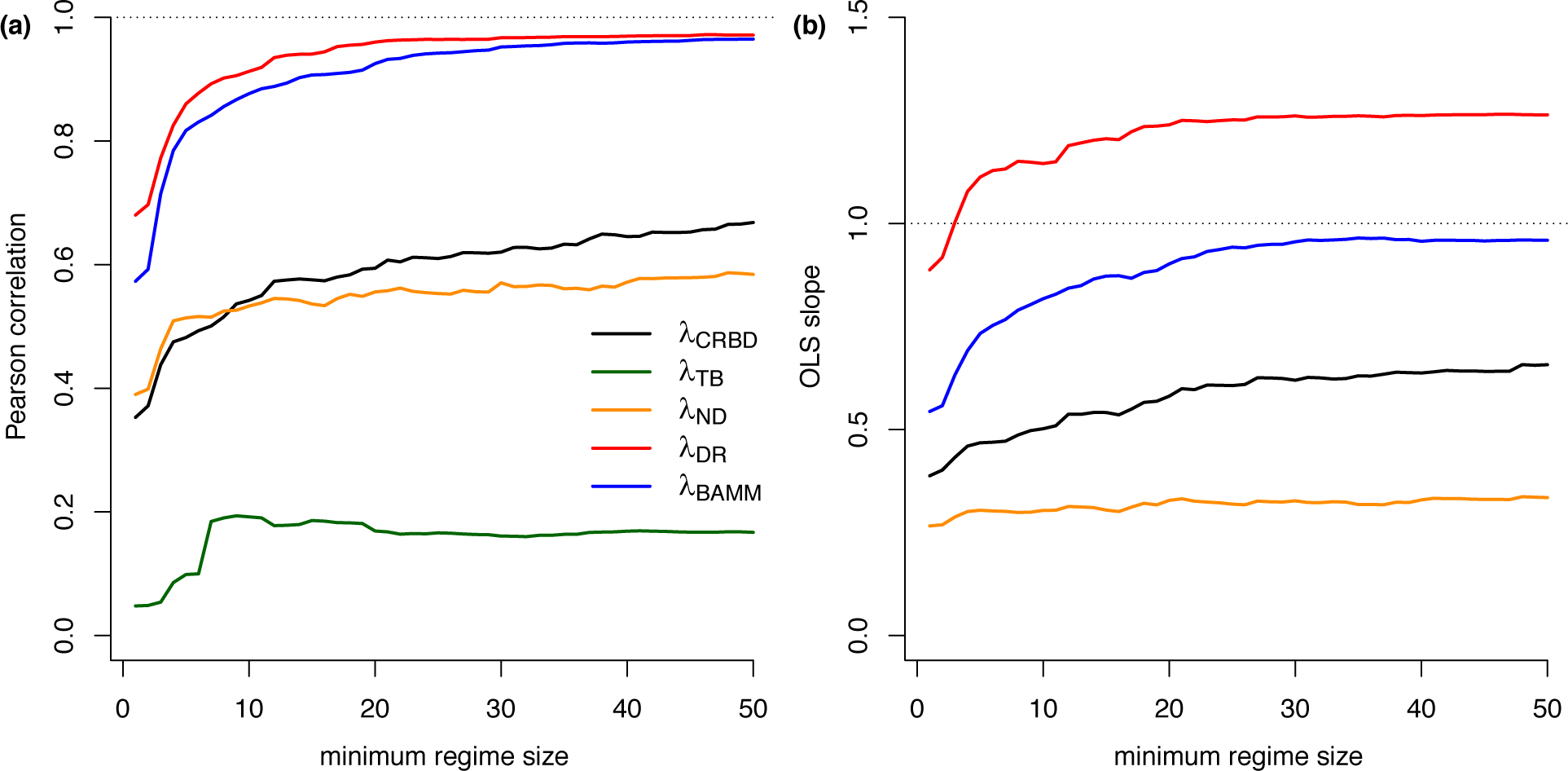
Performance of tip rate metrics as a function of regime size, including Pearson correlation (a) and OLS regression slope (b) for mean rates with respect to λ_TRUE_. λ_DR_ and λ_BAMM_ outperform the other metrics when summarized in this fashion, although λ_DR_ tends to overestimate the rate of speciation. The x-axis denotes the minimum regime size across which performance was summarized. For example, x = 20 corresponds to the correlations and slopes computed for all regimes with 20 or more tips; a value of x = 1 is the corresponding results for all regimes. The OLS slope for λ_TB_ is not visible as it ranges between 7 and 9.

Absolute error in regime mean tip rates was lowest for λ_DR_ and λ_BAMM_, regardless of the size of the rate regime (Figure 5). BAMM’s ability to accurately estimate tip rates improved with regime size, whereas absolute error was relatively consistent across regime sizes for λ_DR_ for regimes greater than 10 species. We also found that λ_DR_ slightly outperformed λ_BAMM_ for small rate regimes.

**Figure 5.**
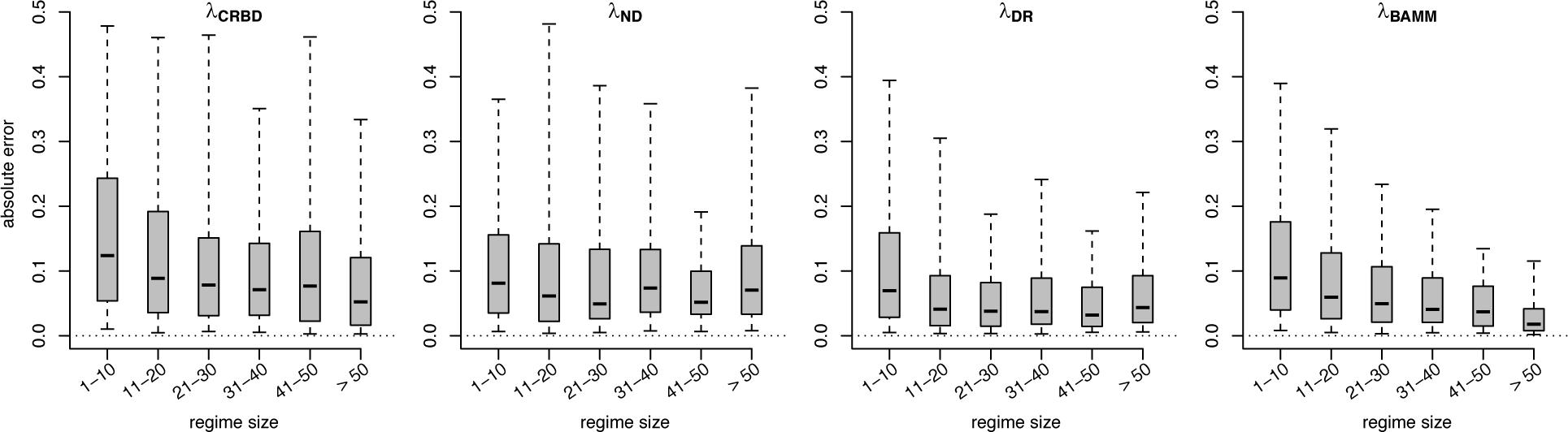
Mean per-regime absolute error in relation to true rate regime size, as binned into 10 size categories. The boxes and whiskers represent the 0.25 – 0.75, and the 0.05 – 0.95 quantile ranges, respectively. Perfectly estimated rates have an error of zero. λ_DR_ and λ_BAMM_ exhibit the least error when averaged by regimes, and λ_DR_ does slightly better for small clades (10-clade median error 0.07 for λ_DR_, and 0.08 for λ_BAMM_).

Note that, in Figures 4 and 5, each rate regime is treated as a single data point. Rate regimes of sizes 1000, 100, and 1 tip are equivalent under this method of error assessment. Figure 4 assesses how well these methods estimate rates for individual regimes, regardless of the size of those regimes. In contrast, Figures 1-3 ask how well these methods perform at estimating rates for a given tip.

## Discussion

We assessed several tip rate metrics and confirmed that these are more accurate estimators of the rate of speciation, rather than net diversification (Figures 1, 4, S7, S10). This distinction was especially pronounced at high extinction fractions, where the rate of lineage turnover is high, and rates of speciation and net diversification have the potential to be more divergent. These results are consistent with supplemental analyses performed in Jetz et al. 2012. It is also important to note that recent extinction will have a much greater influence on these metrics than extinction events deeper in time (Quental & Marshall 2011). Net diversification rate is a critical determinant of species richness, yet this quantity is potentially independent of the underlying rate of speciation. Misinterpretation of tip rate metrics could therefore lead to highly misleading perspectives on large-scale diversity dynamics. As we demonstrate (Figures 1, S2, S3), tip rate metrics (λ_ND_, λ_DR_) provide relatively little information about net diversification, and high values of these metrics are fully consistent with equilibrial models of speciation where the true net diversification rate is zero. Thus, λ_DR_ and λ_ND_ should not be used to support claims about the dynamics of species richness or net diversification *per se* without independent evidence bearing on plausible levels of extinction.

In terms of accuracy, we found that BAMM performed better than non-model-based metrics across all datasets we considered: estimated tip rates were most highly correlated with true tip rates, and mean per-tip error in rates was lower across a range of rate-variable simulation scenarios. This performance is likely to be at least partially due to the inclusion of extinction in the BAMM inference model. BAMM is expected to perform well for phylogenies with discrete shifts in diversification rates as this type of rate variation is most consistent with BAMM’s assumptions (Rabosky 2014; Mitchell & Rabosky 2016; Rabosky, Mitchell & Chang 2017; Mitchell, Etienne & Rabosky 2018). However, BAMM performed surprisingly well for the evolving rates phylogenies, which conform poorly to the assumptions of the inference model. In these trees, the rate of speciation changes continuously under a diffusion process, and as a result, the phylogeny exhibits rate heterogeneity without discrete rate shifts.

On evolving rates phylogenies, λ_BAMM_ performed better than λ_DR_ (Figure 2; Spearman’s ρ for λ_BAMM_ = 0.83, ρ for λ_DR_ = 0.62), despite the fact that λ_DR_ does not rely on the detection of distinct rate regimes to estimate tip rates (Figure 5). λ_BAMM_ also exhibited the lowest mean per-tip error across varying levels of rate heterogeneity (Figure 3).

Why do λ_BAMM_ and λ_DR_ exhibit such striking differences in performance across the simulation scenarios considered here? To illustrate the differences between inference under these metrics, we compared true tip rates to λ_BAMM_ and to λ_DR_ on a simulated birth-death tree with a single rate shift (Figure 6), as well as on one evolving rates tree simulated for this study (Figure 7). It is clear that if BAMM has the statistical power to detect true rate shifts, then it will perform well under rate shift scenarios. In contrast, λ_DR_ tracks true rate shifts but exhibits high sample variance. With an evolving rates tree (Figure 7), the simulation model is very different from the inference model in BAMM. However, it conservatively places rate shifts in order to accommodate rate heterogeneity that is spread across the phylogeny under a rather different model of rate variation. λ_DR_ also broadly tracks the overall pattern of the true rates, but the variance in the corresponding estimates is so high that performance is negatively affected. If we calculate mean (absolute) per-tip error in λ_BAMM_ and λ_DR,_ the error is relatively similar between λ_BAMM_ and λ_DR_, but the variance in per-tip error for λ_DR_ is higher. Overall, BAMM exhibited substantially lower error than λ_DR_ under precisely this scenario (Figure 3).

**Figure 6.**
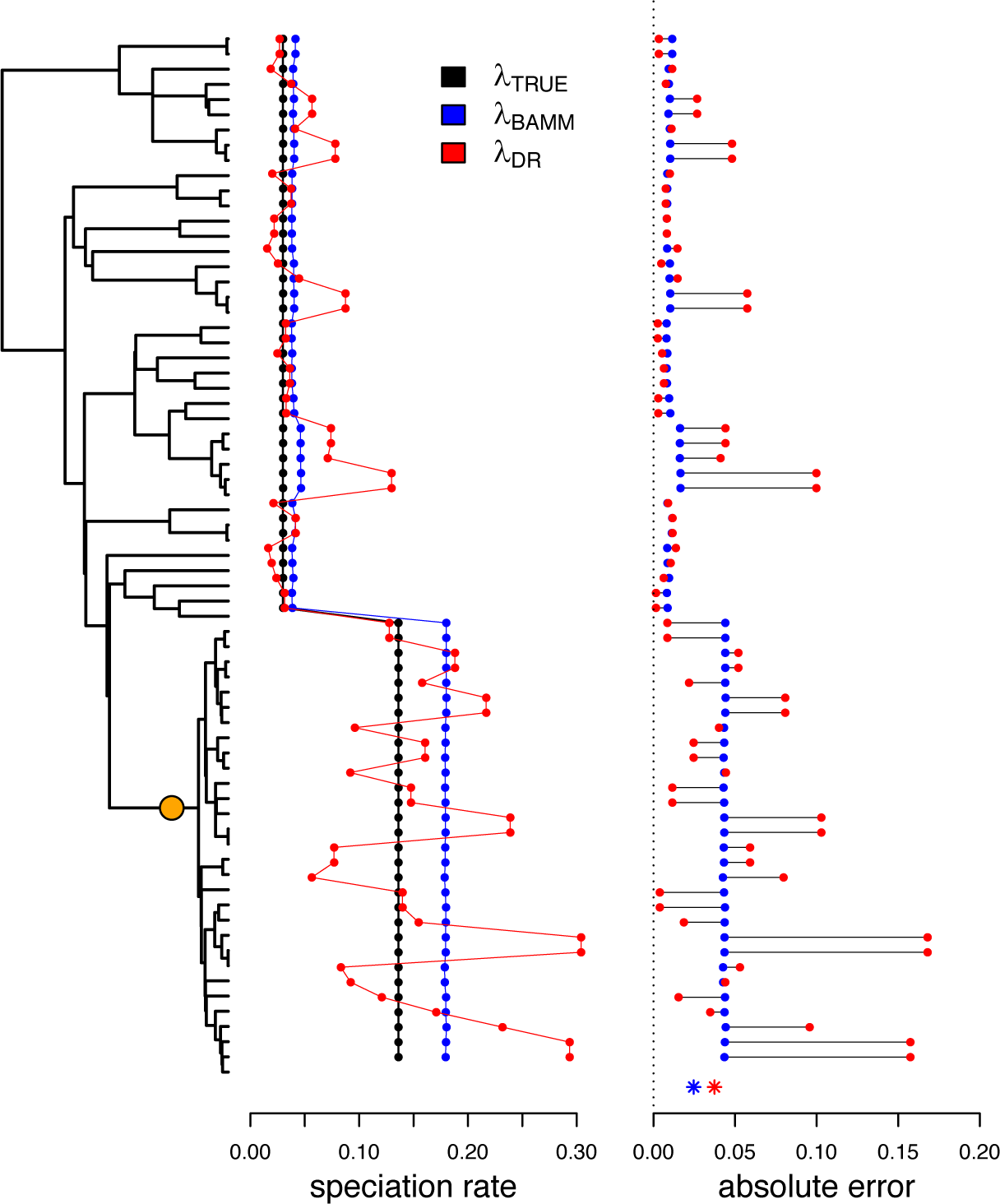
Relationship between λ_TRUE_, λ_BAMM_, and λ_DR_ for a simulated phylogeny containing a single rate shift (orange circle). Subplots to the right of the tree illustrate true and estimated rates for each tip (left) and corresponding absolute error (right). Asterisks at the bottom denote mean per-tip error in tip rate metrics. Mean per-tip error is relatively low and similar between λ_DR_ and λ_BAMM_, but the sample variance in λ_DR_ tip rates is high. In this example, the variance in absolute per-tip error in λ_DR_ is 0.002 versus 0.0003 for λ_BAMM_. On average, λ_DR_ tends to either overestimate or underestimate rates relative to λ_BAMM_, even if the mean per-tip error is relatively low for both metrics.

**Figure 7.**
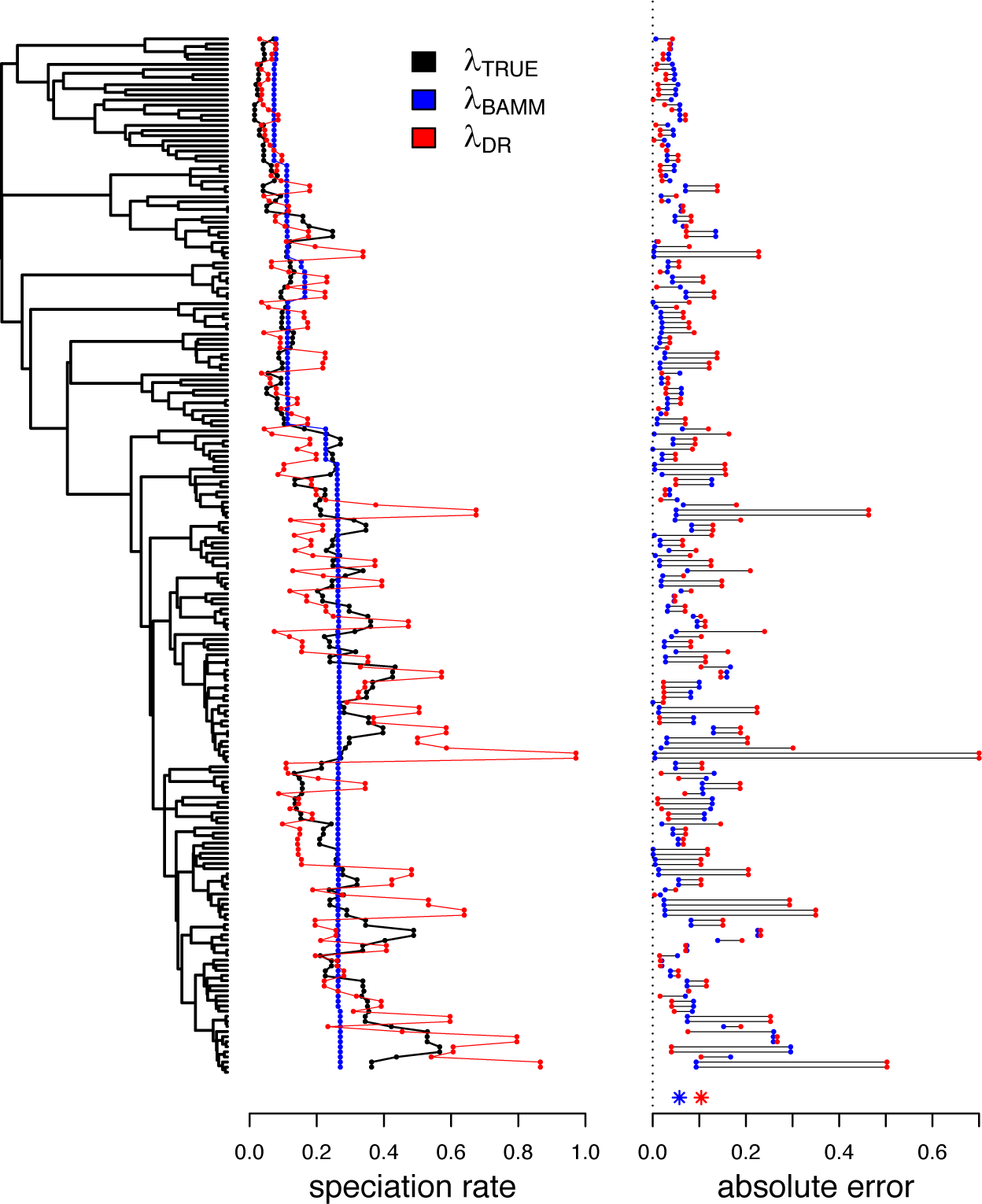
Relationship between λ_DR_, λ_BAMM_, and λ_DR_ for a phylogeny simulated under an “evolving rates” model, such that the speciation rate itself varies under a diffusion model. See Figure 6 for additional details. Neither metric is particularly well equipped to infer the true rate variation in this case. However λ_BAMM_’s conservative estimates are still more accurate relative to λ_DR_, which is negatively impacted by high variance in tip rates. Here, variance in absolute per-tip error in λ_DR_ is 0.012 versus 0.003 for λ_BAMM_.

Thus, although BAMM is conservative in the estimation of tip rates relative to λ_DR_, the method exhibits lower overall error. It appears that λ_DR_ can recover more subtle rate heterogeneity relative to BAMM (see Rabosky, Mitchell & Chang 2017 for discussion of power in BAMM), but this apparent power advantage comes at the cost of increased variance (error) in the resulting estimates. Remarkably, on a per-tip basis, we find that a simple constant-rate birth-death process (λ_CRBD_) frequently yields tip estimates with lower median error and less error variance than those obtained with λ_DR_ (Figure 3), despite the simplifying (and incorrect) assumption that rates are identical across all tips in a given tree. For example, across all multi-regime simulations (Figure 3), λ_CRBD_ point estimates were more accurate than the corresponding λ_DR_ point estimates for 84% of trees in the simulations; for λ_BAMM_, the λ_CRBD_ estimates were more accurate for a much smaller fraction of the total (36%). Similar results were noted for diversity-dependent (λ_CRBD_ more accurate than 98% of λ_DR_ estimates, versus 15% of λ_BAMM_ estimates) and evolving rates trees (λ_CRBD_ more accurate than 93% of λ_DR_ estimates, versus 36% of λ_BAMM_ estimates). Given that λ_DR_ can and does track true heterogeneity in speciation rate (Figures 6, 7), this pattern suggests that the metric is especially sensitive to the stochastic variation in branch lengths that can emerge even when all tips have the same underlying speciation rate.

Regardless of the performance summaries presented in this article, important questions remain with respect to how well tip rate metrics can estimate the true rate of speciation from empirical phylogenies. The phylogenies analyzed in this study were simulated under idealized processes and neglect potential biases and sources of uncertainty that are present in real datasets. For example, if the process of speciation takes time to complete, as is generally believed to be the case (i.e., the protracted speciation process; Rosindell et al. 2010; Etienne and Rosindell 2012), then the most recent speciation events may still be on-going at the present and typical species-level molecular phylogenies may fail to recognize these events. This will lead to an overestimation of terminal branch lengths, as some terminal branches potentially include incipient species. A related bias might arise due to incomplete taxon sampling, which disproportionately affects the length of terminal (or otherwise recent) branch lengths (Pybus & Harvey 2000). Likewise, variation in taxonomic practice across a phylogeny might lead to spurious rate variation, particularly if different species concepts are used, or if some clades in the phylogeny – but not others – have been subject to population genetic analysis or screens for cryptic species diversity. Additionally, it has been shown that BAMM and other methods may fail to infer accurate speciation rate dynamics if the phylogeny is in diversity decline – that is, when extinction rates increase towards the present and ultimately exceed speciation rates (Quental & Marshall 2011; Burin et al. 2018). A major, if obvious, caveat in the interpretation of tip rates is that they apply to recent speciation rates and are necessarily limited with respect to inferences about historical variation in speciation rate.

The greater the importance of the terminal branches in tip rate metrics, the greater the impact these biases might have on tip rate estimates. On one end of the spectrum, metrics such as λ_TB_ will be very sensitive to such biases as they rely exclusively on terminal branch lengths. Such approaches may retain utility as summary statistics (e.g., Bromham, Hua & Cardillo 2016), but we found that λ_TB_ exhibited the greatest amount of error in estimating speciation rates. On the other end of the spectrum, a metric like λ_ND_ would be minimally impacted as this metric is attempting to capture an average speciation rate over an entire root-to-tip path and does not upweight the contribution of recent branch lengths. λ_DR_ is likely somewhere in the middle of this spectrum, as it gives decreasing weight to branches towards the root. λ_BAMM_ is potentially sensitive to such issues as well, although it may be possible to analytically correct for some biases in the mechanics of the model itself (e.g., Rosindell et al. 2010; Etienne and Rosindell 2012).

Potential empirical biases aside, tip rates present a number of practical advantages in the study of diversification rate variation. First, tip rates can be summarized and compared across non-monophyletic assemblages of species (Jetz et al. 2012; Kennedy et al. 2016; Belmaker & Jetz 2015; Oliveira et al. 2016; Quintero & Jetz 2018; Rabosky et al. 2018), making it possible to summarize rate characteristics of entire communities or regional assemblages of species. Second, estimation of rates at the present should be more robust to the influence of extinction, as extinction can erase the history of lineage splitting deeper in the phylogeny (Nee et al. 1994; Nee, May & Harvey 1994; Rabosky & Lovette 2008). Third, tip-specific rates can be paired with species-specific trait values or geographic attributes in order to test potential trait-or geography-dependent speciation rates (Freckleton, Phillimore & Pagel 2008; Jetz et al. 2012, Rabosky & Goldberg 2017; Harvey & Rabosky 2017). Tip rates make it possible to relax strong assumptions of rate homogeneity within character states, which are inherent to certain trait-dependent models, including BiSSE and GeoSSE (Maddison, Midford & Otto 2007; Goldberg, Lancaster & Ree 2011; Ng & Smith 2014). Recent work has provided a conceptually rich and robust interpretive framework for SSE models that does not assume rate-constancy within character states (Beaulieu & O’Meara 2016; Caetano, O’Meara & Beaulieu 2018), but tip rates nonetheless can provide an important check on results obtained with SSE models by providing a direct means of visualizing the relationship between branch lengths and character states (Bromham, Hua & Cardillo 2016; Hua & Bromham 2016; Harvey & Rabosky 2017). Visual inspection of data in this fashion has the potential to reduce false positives by calling attention to potential outliers and other sources of model inadequacy (Maddison & FitzJohn 2014; Rabosky & Goldberg 2015). A final advantage for non-model-based tip rates, especially λ_DR_, is that they can profitably be applied to extremely large phylogenies: there are few computational limits to using them on phylogenies with tens of thousands of tips or more, in contrast to formal model-based approaches for which BAMM, HiSSE (Hidden State Speciation and Extinction; Beaulieu & O’Meara 2016), and other methods are poorly suited. This computational efficiency also lends itself to more readily accounting for phylogenetic uncertainty, because tip rate metrics can rapidly be computed across posterior distributions of phylogenies and averaged (for example, see Jetz et al. 2012; Rabosky et al. 2018).

In summary, tip rates offer a number of theoretical and practical advantages, particularly in the study of associations between traits and diversification. We found that λ_BAMM_ outperformed other metrics evaluated in this study and proved to be relatively accurate, even under diversification scenarios that depart from the BAMM inference model. λ_DR_ underperformed in comparison to λ_BAMM_, but in many cases still did reasonably well, particularly for small rate regimes. Despite our performance results, λ_DR_ is likely to remain a useful tool in the study of trait-and geography-dependent diversification (Rabosky & Goldberg 2017; Harvey & Rabosky 2017).

## Acknowledgements

We thank Jonathan Mitchell for help with compiling the simulation datasets evaluated in this study. We thank Francisco Henao Diaz, Matt Pennell, Brian O’Meara, the O’Meara Lab, and an anonymous reviewer for comments on the manuscript. We also thank Michael Grundler, as well as all other members of the Rabosky and Davis Rabosky labs at the University of Michigan for thoughtful discussion. This work was supported in part by a University of Michigan Rackham Predoctoral Fellowship (P.O.T.) and by a Fellowship from the David and Lucile Packard Foundation (D.L.R.).

## Data accessibility

All data and code necessary to repeat the analyses have been made available through the Dryad digital data repository at XXXXXX.

## Author contributions

P.O.T. and D.L.R. designed the project. P.O.T. assembled the datasets and performed all analyses. P.O.T. and D.L.R. wrote the manuscript. Both authors contributed critically to subsequent drafts and approved the final publication.

---

**Figure S1.**
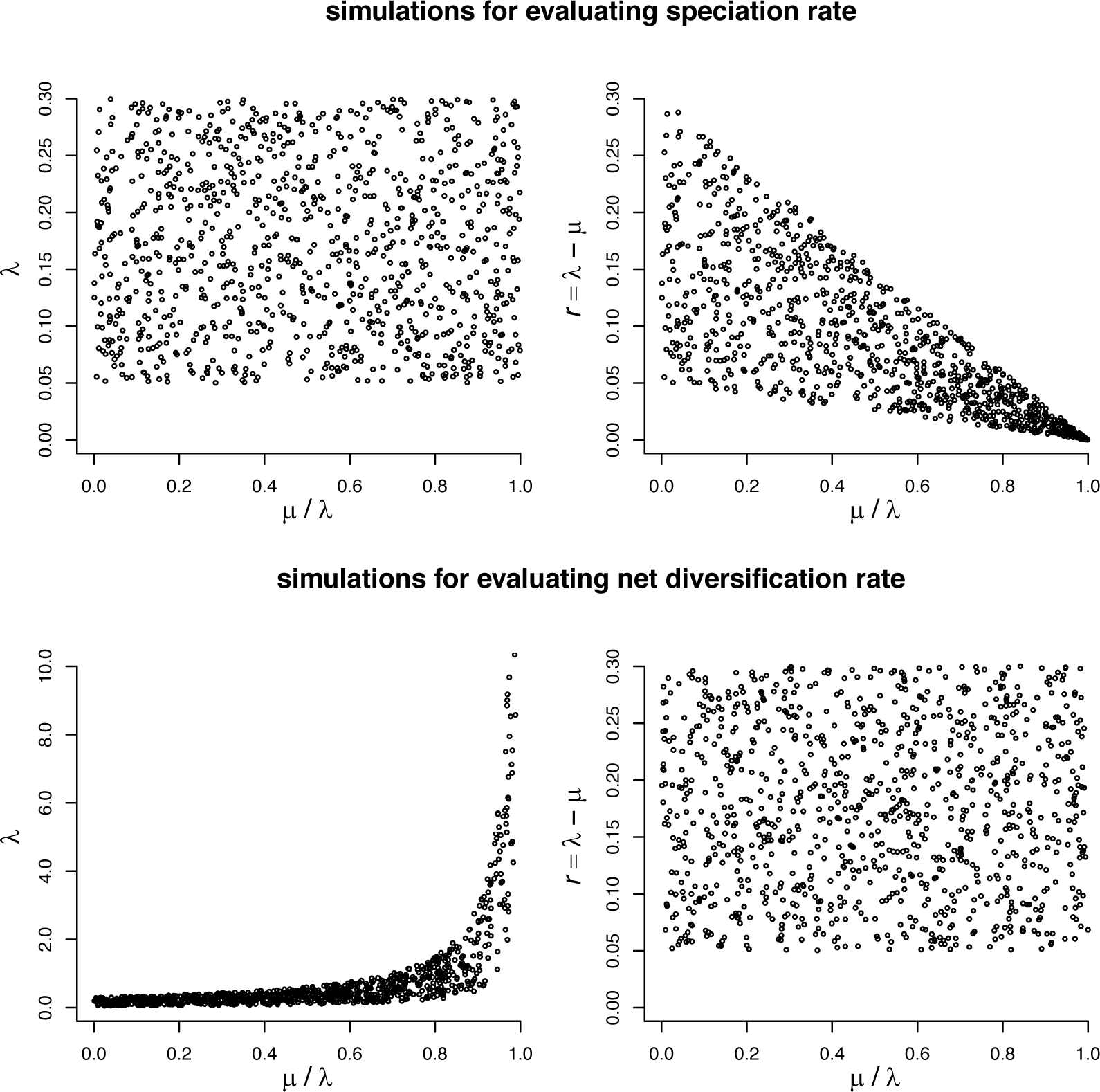
Details of the phylogeny simulations designed to evaluate the performance of the four tip metrics in terms of speciation rate and net diversification rate. From the top row, it is clear that when λ is sampled uniformly with respect to ε, the distribution of *r* is not uniform: the mean, range and variance in *r* decrease dramatically as ε increases. The reverse is true for the distribution of λ when *r* is sampled uniformly with respect to ε (bottom row). Our simulation design ensures that λ and *r* are sampled from identical uniform distributions with respect to ε and ensures comparability of the resulting error estimates.

**Figure S2.**
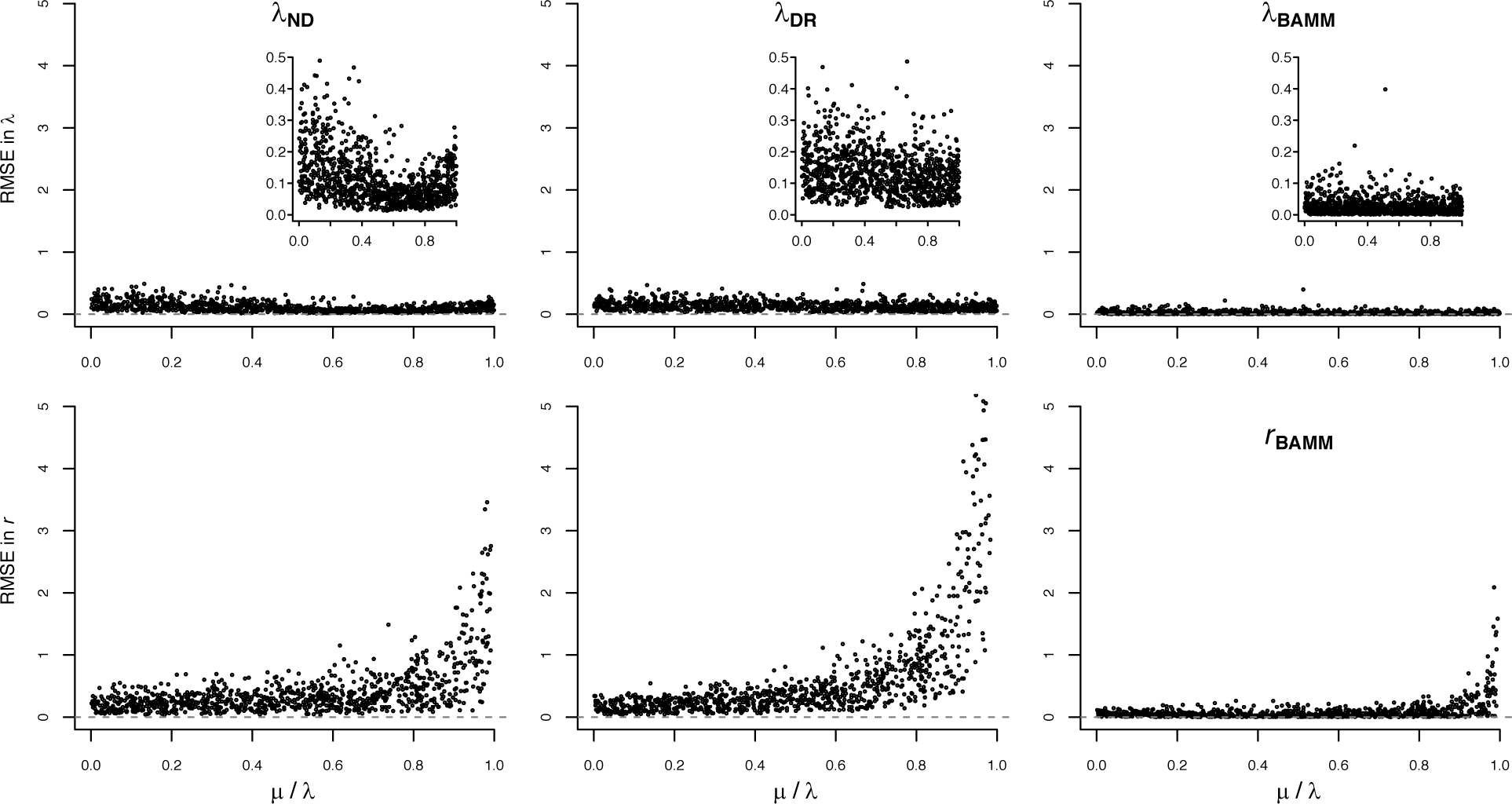
Root-mean-square error (RMSE) in λ (top) and *r* (bottom) for three different tip rate metrics, across a range of relative extinction rates. For BAMM, the estimated speciation and net diversification rates are presented in the top and bottom panels, respectively. Error of zero implies perfect accuracy. Inset plots show error in λ with truncated y-axis scale to facilitate comparison among metrics. All tip rate metrics track λ more accurately than they track *r*.

**Figure S3.**
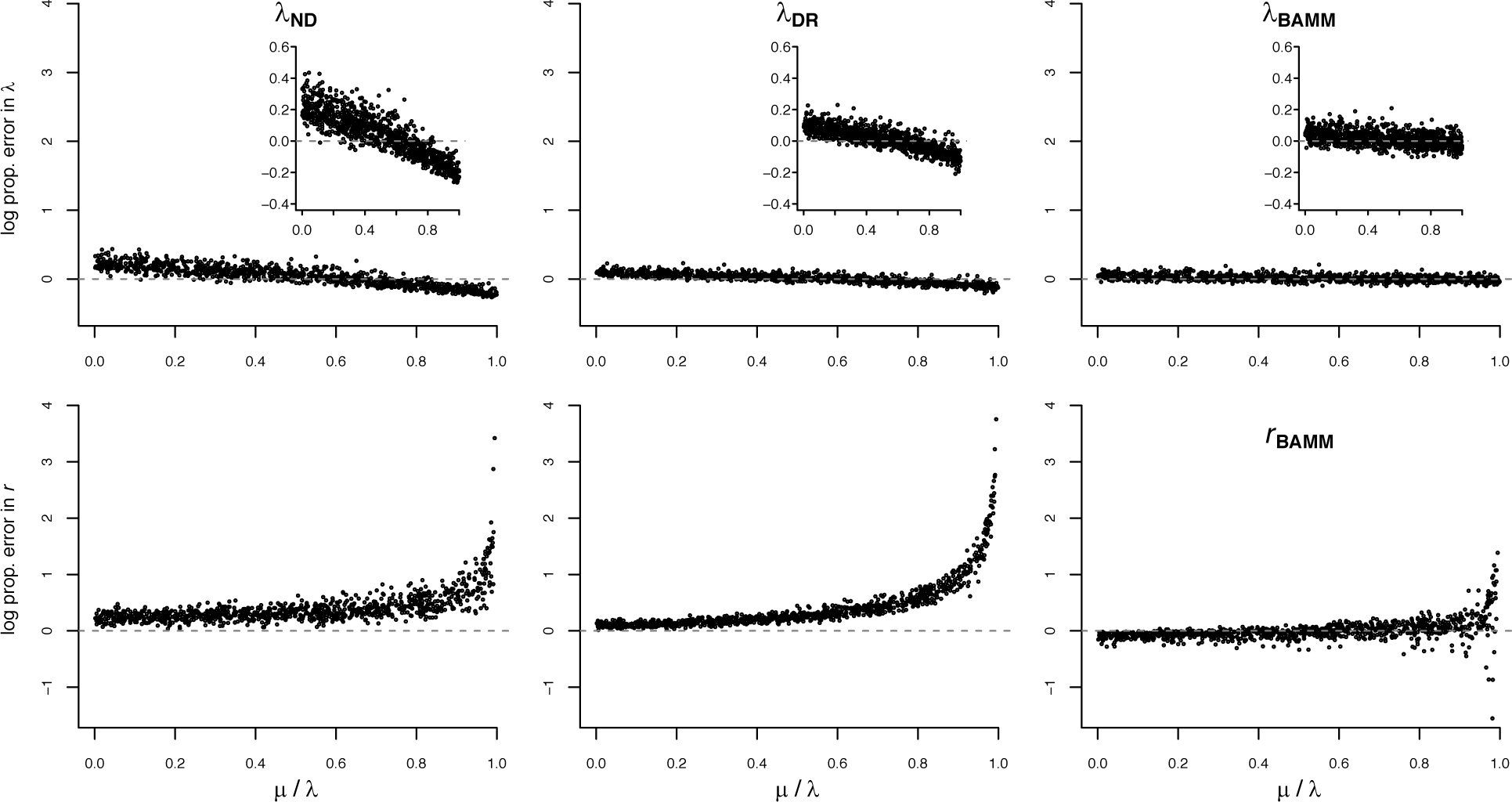
Log proportional accuracy in λ (top) and *r* (bottom) for different tip rate metrics, across a range of relative extinction rates. For BAMM, the estimated net diversification rate is presented. Proportional error of 0 implies perfect accuracy. Inset plots reveal greater detail in error for λ to ease metric comparison. All tip metrics track λ much more accurately than they track *r*, and λ_BAMM_ does so with the least amount of error. See Figure S4 for λ_TB_.

**Figure S4.**
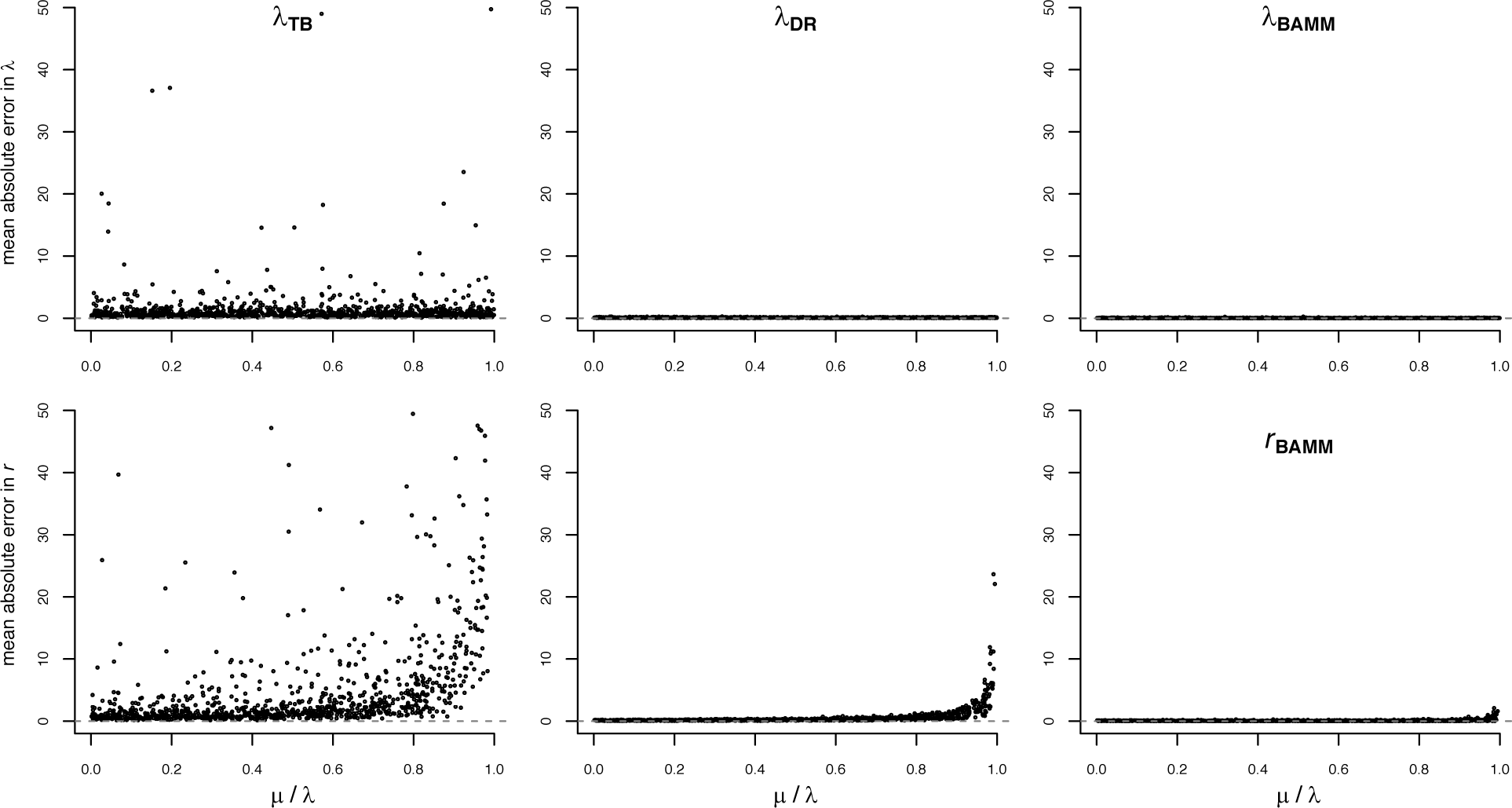
Mean absolute error in λ (top) and *r* (bottom) for λ_TB_, with λ_DR_ and λ_BAMM_ on the same scale for comparison. For BAMM, the estimated net diversification rate is presented. λ_TB_ more accurately tracks λ and than *r*, but the amount of error is an order of magnitude greater than for other metrics.

**Figure S5.**
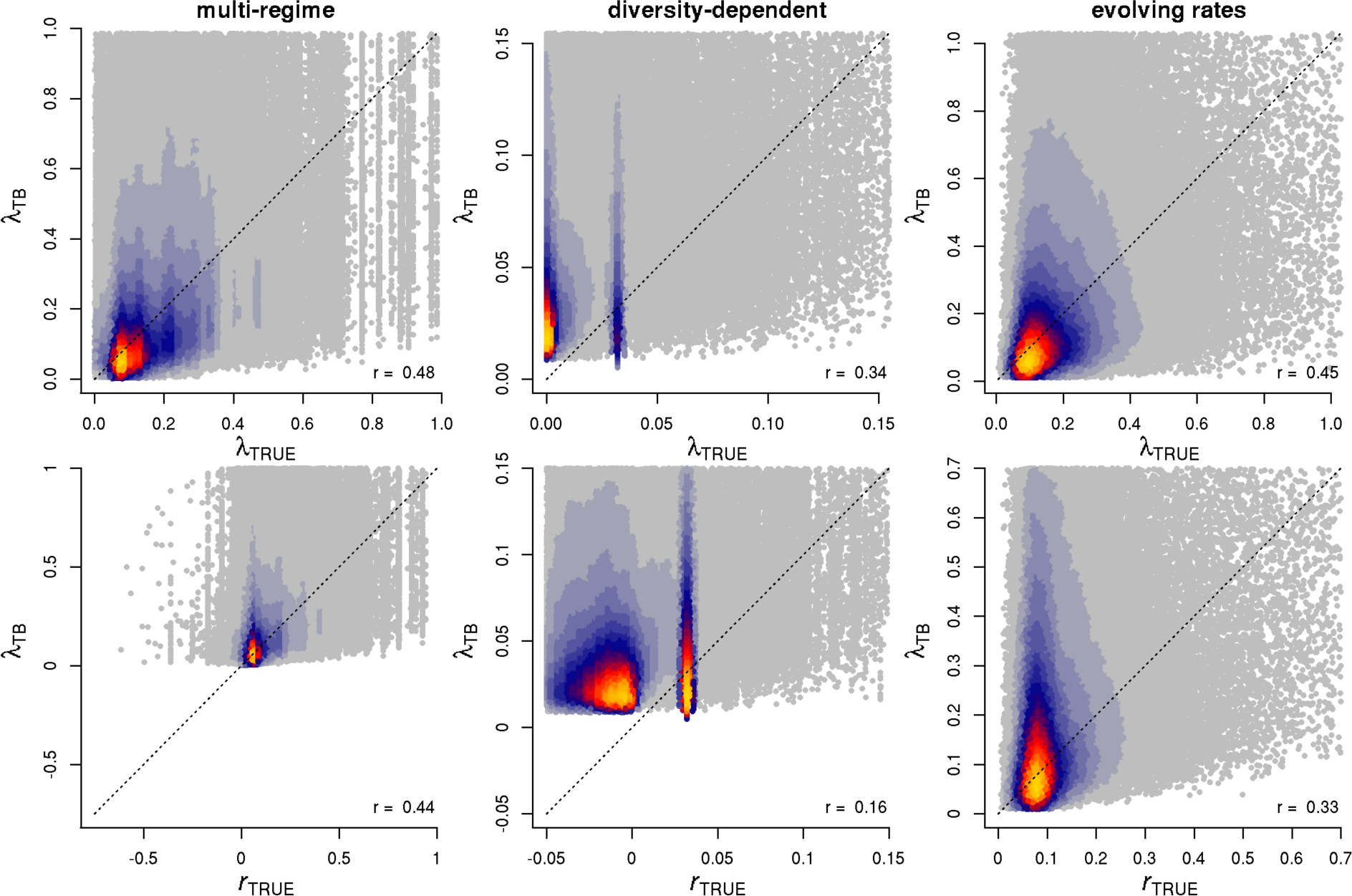
True tip rates (top row: λ_TRUE_, bottom row: *r*_TRUE_) in relation to λ_TB_. Tip rates were compared separately for different major categories of phylogeny simulations (rows). Plotting region is restricted to the 99th percentile of true rates, but Spearman correlations between true and estimated rates (lower right of each figure panel) are based on the full range of the data. Colors indicate the density of points in the scatter plots. λ_TB_ is a largely unbiased but noisy measure of true speciation rate.

**Figure S6.**
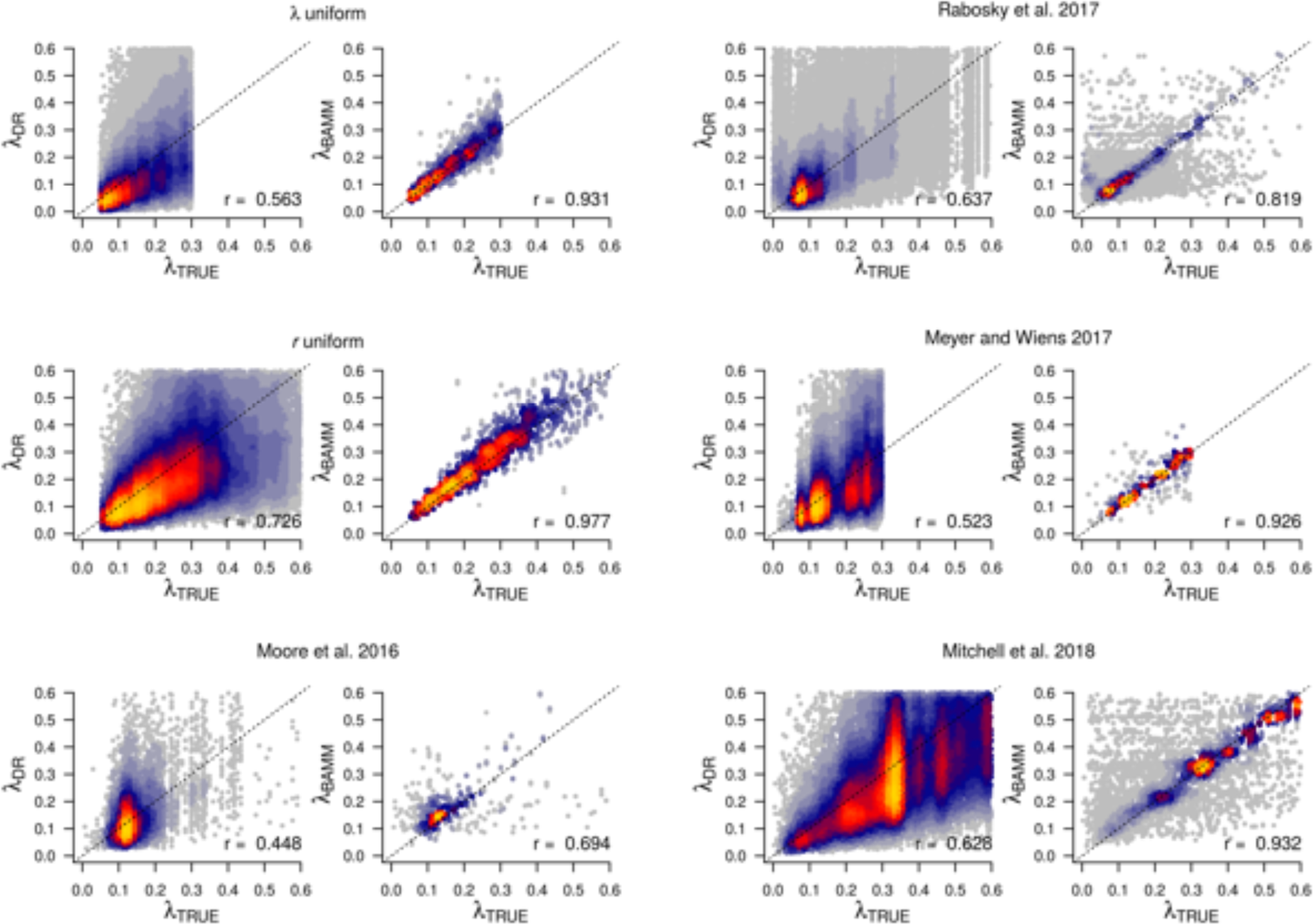
True tip rates (λ_TRUE_) in relation to estimated tip rates from multi-regime phylogenies, as inferred from the best two tip rate metrics, λ_DR_ and λ_BAMM_. Data are separated by source, to confirm that patterns described in the main text are not driven by any one simulation study. Spearman’s correlation is presented in the bottom right corner. Colors indicate the density of points in the scatter plots. Regardless of the dataset, λ_BAMM_ performs noticeably better than λ_DR_.

**Figure S7.**
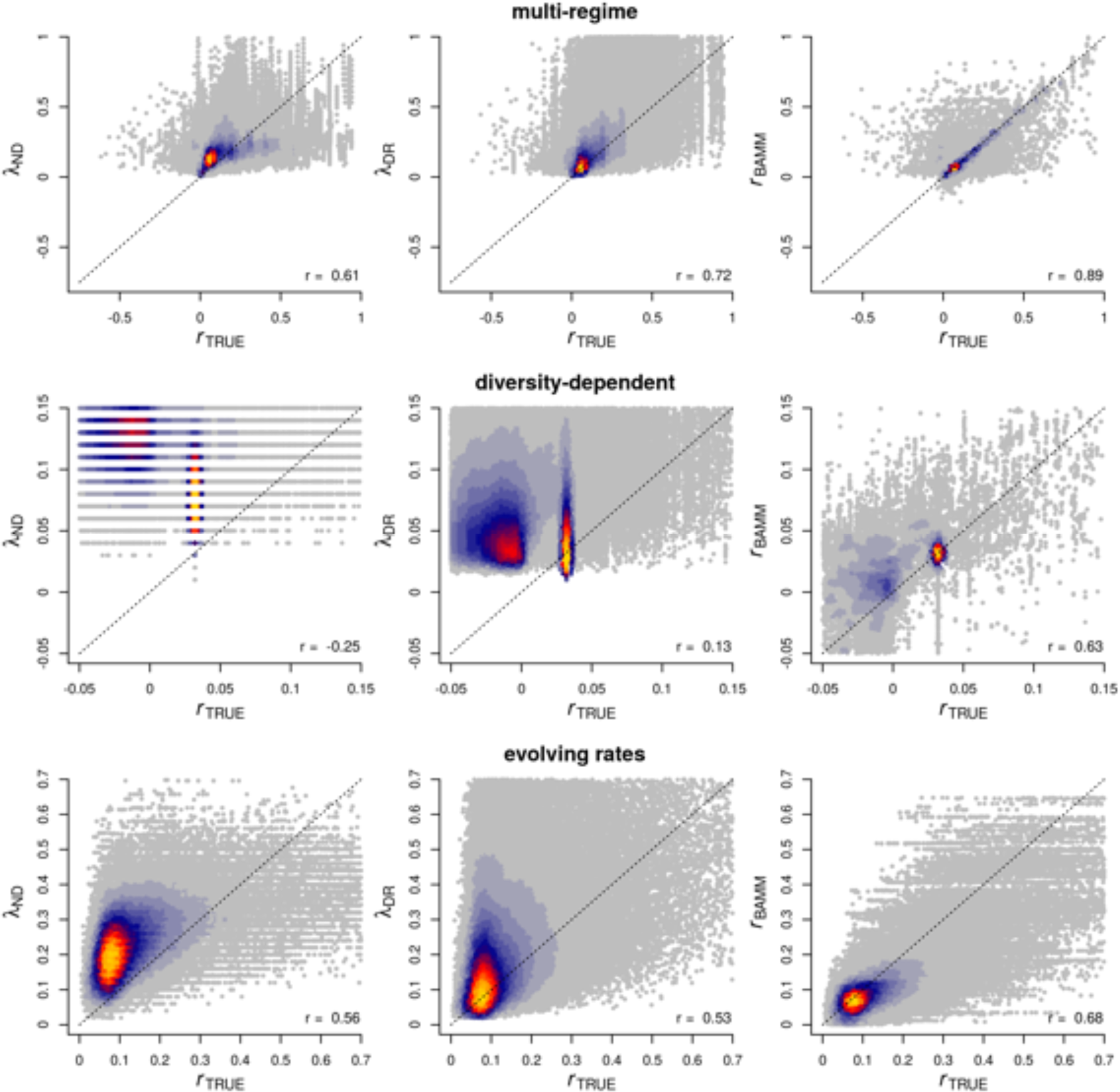
True net diversification tip rates (*r*_TRUE_) in relation to estimated tip rates. Tip rates were compared separately for different major categories of phylogeny simulations (rows). Plotting region is restricted to the 99th percentile of true rates, but Spearman correlations between true and estimated rates (lower right of each figure panel) are based on the full range of the data. Colors indicate the density of points in the scatter plots. The horizontal gaps in λ_ND_ for diversity-dependent trees are an artefact of all trees having the same crown age. Relative performance comparison aside, correlations with *r*_TRUE_ are lower than with λ_TRUE_ (Figure 2).

**Figure S8.**
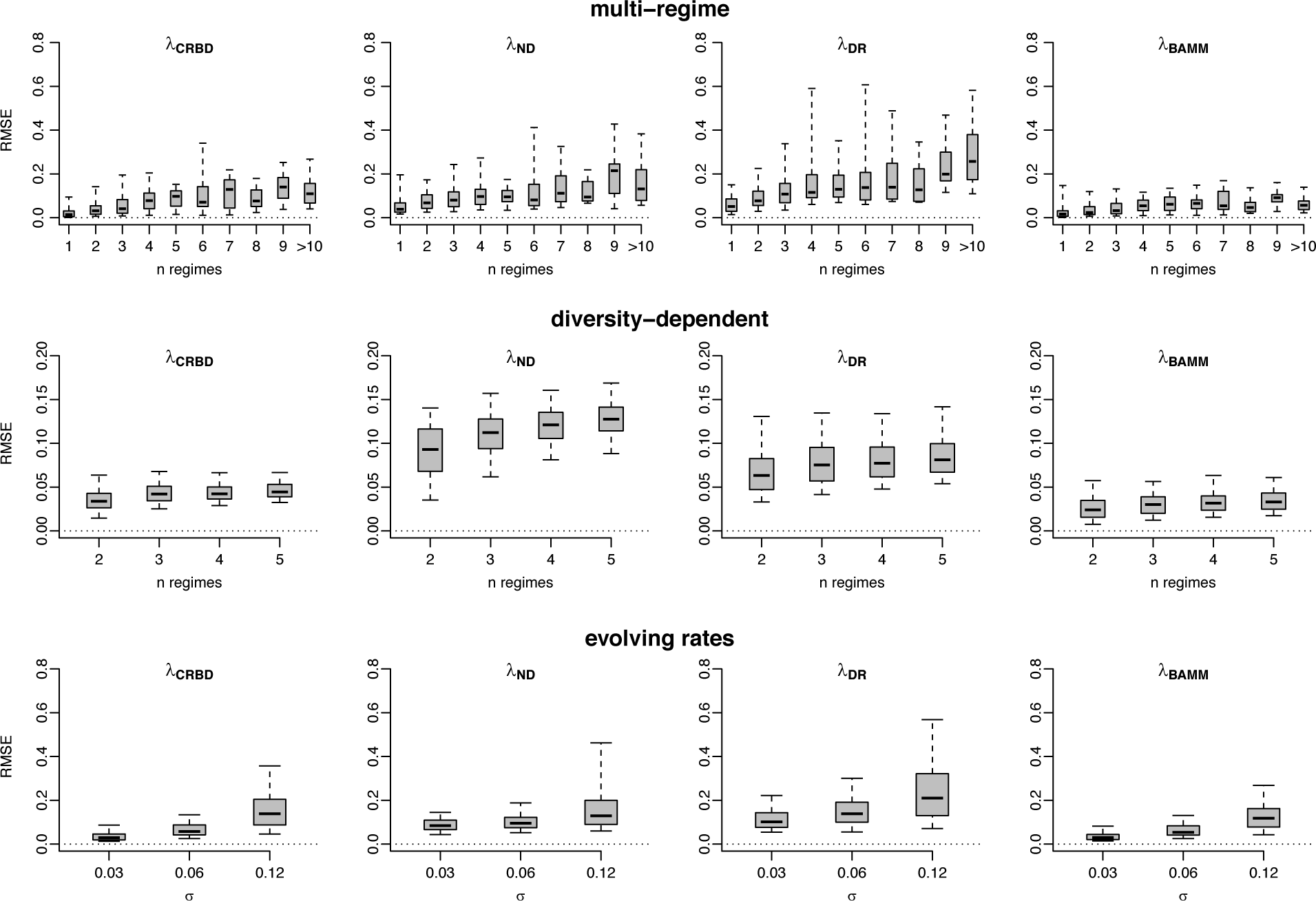
Root-mean-square error (RMSE) in speciation rates as a function of the magnitude of rate heterogeneity in each simulated phylogeny. Results are presented separately for different categories of rate variation (Table 1); left column shows estimates from a constant-rate birth-death model for reference. The boxes and whiskers represent the 0.25 – 0.75, and the 0.05 – 0.95 quantile ranges, respectively. In some cases, λ_ND_ and λ_DR_ had more error than a simple CRBD model with no variation in tip rates. λ_BAMM_ had the least amount of error across all amounts of rate heterogeneity.

**Figure S9.**
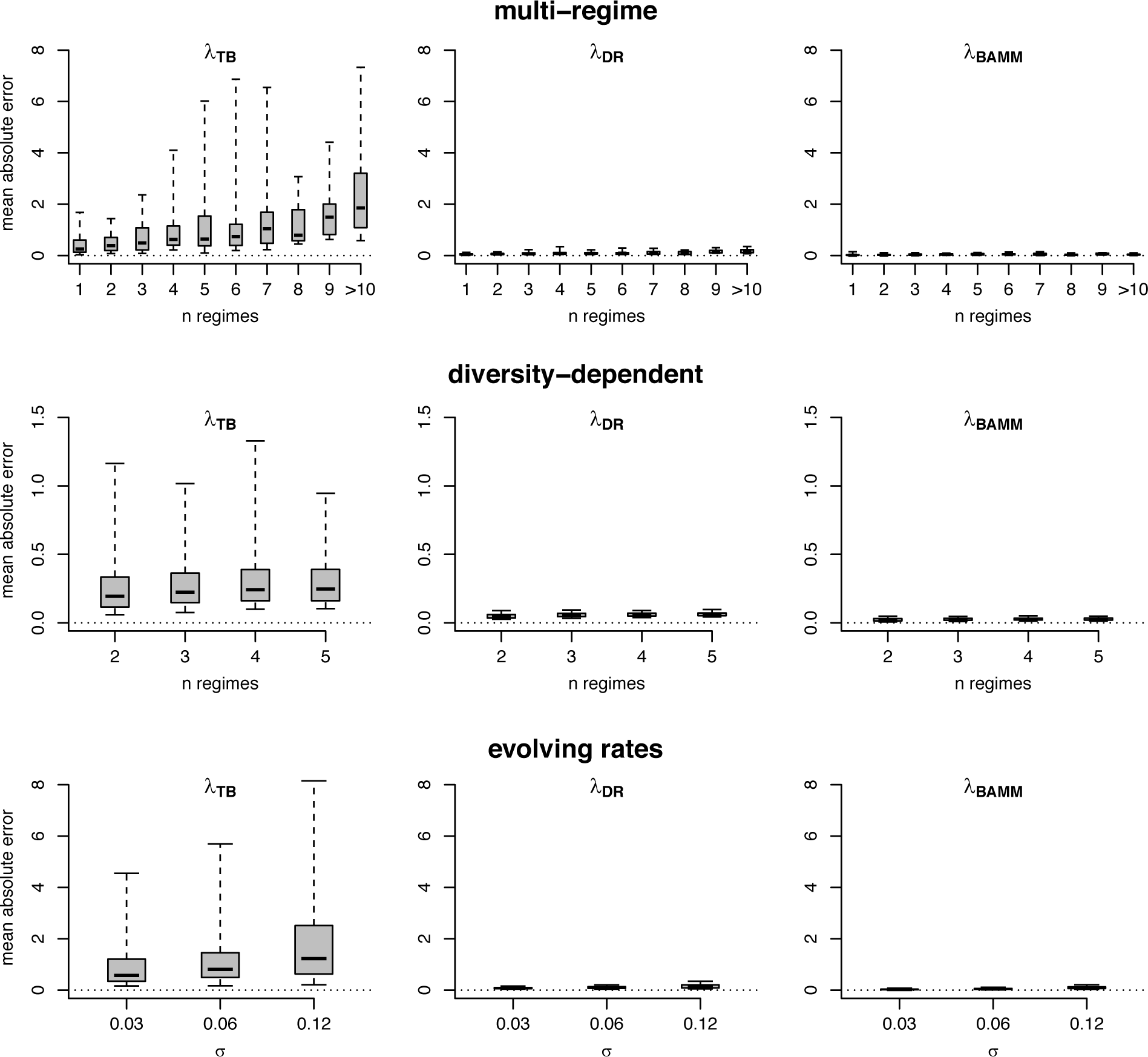
Mean per-tip absolute error in λ_TB_ as a function of the magnitude of rate heterogeneity in each simulated phylogeny. λ_DR_ and λ_BAMM_ are included on the same scale for comparison. Results are presented separately for different categories of rate variation (Table 1). The boxes and whiskers represent the 0.25 – 0.75, and the 0.05 – 0.95 quantile ranges, respectively. Error in λ_TB_ generally increases with increasing rate heterogeneity, and this error is substantially greater than error in other tip rate metrics.

**Figure S10.**
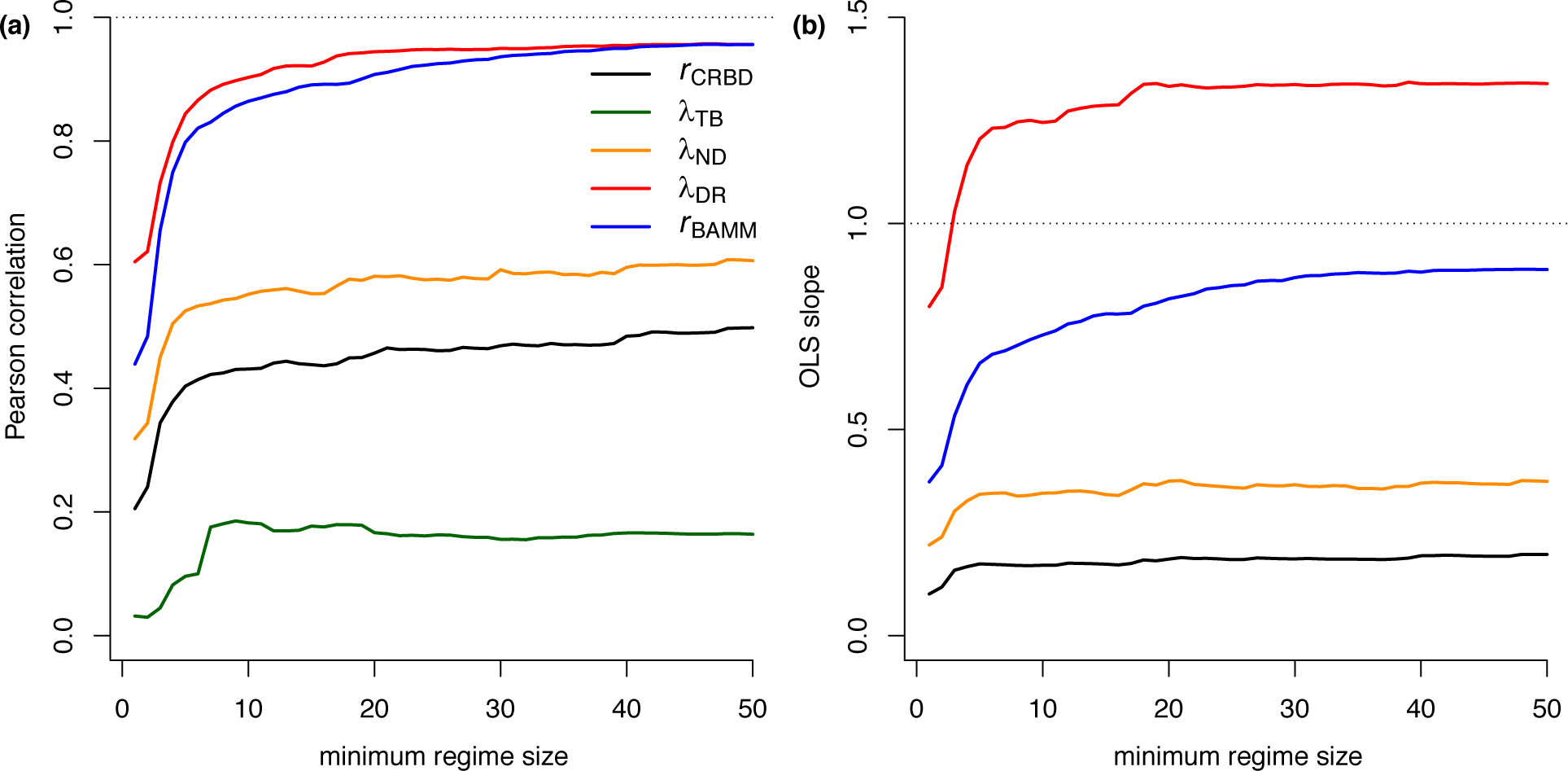
Performance of tip rate metrics as a function of regime size, including Pearson correlation (a) and OLS regression slope (b) for mean rates with respect to *r*_TRUE_. λ_DR_ and *r*_BAMM_ outperform the other metrics when summarized in this fashion, although λ_DR_ overestimates the rate of net diversification (more so than it overestimated λ_TRUE_, Figure 4). The x-axis denotes the minimum regime size across which performance was summarized. For example, x = 20 corresponds to the correlations and slopes computed for all regimes with 20 or more tips; a value of x = 1 is the corresponding results for all regimes. The OLS slope for λ_TB_ is not visible as it ranges between 7 and 9.

## References

Alfaro, M.E., Faircloth, B.C., Harrington, R.C., Sorenson, L., Friedman, M., Thacker, C.E., Oliveros, C.H., Černý, D. & Near, T.J. (2018). Explosive diversification of marine fishes at the Cretaceous–Palaeogene boundary. Nature Ecology & Evolution, 2, 688–696.

Alfaro, M.E., Santini, F., Brock, C., Alamillo, H., Dornburg, A., Rabosky, D.L., Carnevale, G. & Harmon, L.J. (2009). Nine exceptional radiations plus high turnover explain species diversity in jawed vertebrates. Proceedings of the National Academy of Sciences, 106, 13410–13414.

Barker, F.K., Burns, K.J., Klicka, J., Lanyon, S.M. & Lovette, I.J. (2013). Going to extremes: contrasting rates of diversification in a recent radiation of new world passerine birds. Systematic Biology, 62, 298–320.

Beaulieu, J.M. & O’Meara, B.C. (2016). Detecting Hidden Diversification Shifts in Models of Trait-Dependent Speciation and Extinction. Systematic Biology, 65, 583–601.

Beaulieu, J.M. & O’Meara, B.C. (2015). Extinction can be estimated from moderately sized molecular phylogenies. Evolution, 69, 1036–1043.

Belmaker, J. & Jetz, W. (2015). Relative roles of ecological and energetic constraints, diversification rates and region history on global species richness gradients. Ecology Letters, 18, 563–571.

Bromham, L., Hua, X. & Cardillo, M. (2016). Detecting Macroevolutionary Self-Destruction from Phylogenies. Systematic Biology, 65, 109–127.

Burin, G., Alencar, L.R.V., Chang, J., Alfaro, M.E. & Quental, T.B. (2018). How Well Can We Estimate Diversity Dynamics for Clades in Diversity Decline? Systematic Biology, 105, 1–17.

Caetano, D.S., O’Meara, B.C. & Beaulieu, J.M. (2018). Hidden state models improve state-dependent diversification approaches, including biogeographical models. Evolution.

Cai, T., Fjeldså, J., Wu, Y., Shao, S., Chen, Y., Quan, Q., Li, X., Song, G., Qu, Y., Qiao, G. & Lei, F. (2017). What makes the Sino-Himalayan mountains the major diversity hotspots for pheasants? Journal of Biogeography, 45, 640–651.

Claramunt, S. (2010). Discovering exceptional diversifications at continental scales: the case of the endemic families of Neotropical suboscine passerines. Evolution, 64, 2004–2019.

Coyne, J.A. & Orr, H.A. (2004). Speciation. Sinauer, Cambridge.

Etienne, R.S. & Haegeman, B. (2012). A Conceptual and Statistical Framework for Adaptive Radiations with a Key Role for Diversity Dependence. The American Naturalist, 180, E75–E89.

Etienne, R.S. & Rosindell, J. (2012). Prolonging the Past Counteracts the Pull of the Present: Protracted Speciation Can Explain Observed Slowdowns in Diversification. Systematic Biology, 61, 204–213.

FitzJohn, R.G. (2010). Quantitative traits and diversification. Systematic Biology, 59, 619–633.

FitzJohn, R.G., Maddison, W.P. & Otto, S.P. (2009). Estimating Trait-Dependent Speciation and Extinction Rates from Incompletely Resolved Phylogenies. Systematic Biology, 58, 595–611.

Freckleton, R.P., Phillimore, A.B. & Pagel, M. (2008). Relating Traits to Diversification: A Simple Test. The American Naturalist, 172, 102–115.

Goldberg, E.E., Lancaster, L.T. & Ree, R.H. (2011). Phylogenetic inference of reciprocal effects between geographic range evolution and diversification. Systematic Biology, 60, 451–465.

Gomes, A.C.R., Sorenson, M.D. & Cardoso, G.C. (2016). Speciation is associated with changing ornamentation rather than stronger sexual selection. Evolution, 70, 2823–2838.

Harvey, M.G. & Rabosky, D.L. (2017). Continuous traits and speciation rates: Alternatives to state-dependent diversification models (N. Cooper, Ed.). Methods in Ecology and Evolution, 12, 751–10.

Hua, X. & Bromham, L. (2016). PHYLOMETRICS: an R package for detecting macroevolutionary patterns, using phylogenetic metrics and backward tree simulation. Methods in Ecology and Evolution, 7, 806–810.

Jablonski, D. (2008). Species selection: Theory and data. Annual Review of Ecology, Evolution, and Systematics, 39, 501–524.

Jetz, W., Thomas, G.H., Joy, J.B., Hartmann, K. & Mooers, A.O. (2012). The global diversity of birds in space and time. Nature, 491, 444–448.

Kay, K., Voelckel, C., Yang, J.Y., Hufford, K.M., Kaska, D.D. & Hodges, S.A. (2006). Floral characters and species diversification. Ecology and Evolution of Flowers (eds L. Harder & S. Barrett), pp. 311–325. Oxford.

Kennedy, J.D., Borregaard, M.K., Jønsson, K.A., Holt, B., Fjeldså, J. & Rahbek, C. (2016). Does the colonization of new biogeographic regions influence the diversification and accumulation of clade richness among the Corvides (Aves: Passeriformes)? Evolution, 38–50.

Maddison, W.P. & FitzJohn, R.G. (2014). The Unsolved Challenge to Phylogenetic Correlation Tests for Categorical Characters. Systematic Biology, 64, 127–136.

Maddison, W.P., Midford, P.E. & Otto, S. (2007). Estimating a binary character’s effect on speciation and extinction. Systematic Biology, 56, 701.

Marin, J. & Hedges, S.B. (2016). Time best explains global variation in species richness of amphibians, birds and mammals. Journal of Biogeography, 43, 1069–1079.

Meyer, A.L.S. & Wiens, J.J. (2017). Estimating diversification rates for higher taxa: BAMM can give problematic estimates of rates and rate shifts. Evolution, 72, 39–53.

Mitchell, J.S. & Rabosky, D.L. (2016). Bayesian model selection with BAMM: effects of the model prior on the inferred number of diversification shifts. Methods in Ecology and Evolution, 8, 37–46.

Mitchell, J.S., Etienne, R.S. & Rabosky, D.L. (2018). Inferring Diversification Rate Variation From Phylogenies With Fossils. Systematic Biology.

Mittelbach, G.G., Schemske, D.W., Cornell, H.V., Allen, A.P., Brown, J.M., Bush, M.B., Harrison, S.P., Hurlbert, A.H., Knowlton, N., Lessios, H.A., McCain, C.M., McCune, A.R., McDade, L.A., McPeek, M.A., Near, T.J., Price, T.D., Ricklefs, R.E., Roy, K., Sax, D.F., Schluter, D., Sobel, J.M. & Turelli, M. (2007). Evolution and the latitudinal diversity gradient: speciation, extinction and biogeography. Ecology Letters, 10, 315–331.

Moen, D. & Morlon, H. (2014). Why does diversification slow down? Trends in Ecology and Evolution, 29, 190–197.

Moore, B.R., Höhna, S., May, M.R., Rannala, B. & Huelsenbeck, J.P. (2016). Critically evaluating the theory and performance of Bayesian analysis of macroevolutionary mixtures. Proceedings of the National Academy of Sciences, 113, 9569–9574.

Near, T.J., Dornburg, A., Kuhn, K.L., Eastman, J.T., Pennington, J.N., Patarnello, T., Zane, L., Fernandez, D.A. & Jones, C.D. (2012). Ancient climate change, antifreeze, and the evolutionary diversification of Antarctic fishes. Proceedings of the National Academy of Sciences, 109, 3434–3439.

Nee, S., Holmes, E.C., May, R.M. & Harvey, P.H. (1994a). Extinction Rates Can Be Estimated From Molecular Phylogenies. Philosophical Transactions of the Royal Society B: Biological Sciences, 344, 77–82.

Nee, S., May, R.M. & Harvey, P.H. (1994b). The reconstructed evolutionary process. Philosophical Transactions of the Royal Society B: Biological Sciences, 344, 305–311.

Nee, S., Mooers, A. & Harvey, P.H. (1992). Tempo and Mode of Evolution Revealed From Molecular Phylogenies. Proceedings of the National Academy of Sciences, 89, 8322–8326.

Ng, J. & Smith, S.D. (2014). How traits shape trees: new approaches for detecting character state-dependent lineage diversification. Journal of Evolutionary Biology, 27, 2035–2045.

Oliveira, B.F., Machac, A., Costa, G.C., Brooks, T.M., Davidson, A.D., Rondinini, C. & Graham, C.H. (2016). Species and functional diversity accumulate differently in mammals. Global Ecology and Biogeography, 25, 1119–1130.

Pybus, O.G. & Harvey, P.H. (2000). Testing macro-evolutionary models using incomplete molecular phylogenies. Proceedings of the Royal Society of London. Series B: Biological Sciences, 267, 2267–2272.

Quental, T.B. & Marshall, C.R. (2011). The Molecular Phylogenetic Signature of Clades in Decline. PLoS ONE, 6, e25780–9.

Quintero, I. & Jetz, W. (2018). Global elevational diversity and diversification of birds. Nature, 555, 246–250.

Rabosky, D.L. (2014). Automatic detection of key innovations, rate shifts, and diversity-dependence on phylogenetic trees. PLoS ONE, 9, e89543.

Rabosky, D.L. (2010). Extinction rates should not be estimated from molecular phylogenies. Evolution, 64, 1816–1824.

Rabosky, D.L. & Goldberg, E.E. (2017). FiSSE: A simple nonparametric test for the effects of a binary character on lineage diversification rates. Evolution, 106, 13410–11.

Rabosky, D.L. & Goldberg, E.E. (2015). Model Inadequacy and Mistaken Inferences of Trait-Dependent Speciation. Systematic Biology, 64, 340–355.

Rabosky, D.L. & Lovette, I.J. (2008). Explosive evolutionary radiations: decreasing speciation or increasing extinction through time? Evolution, 62, 1866–1875.

Rabosky, D.L., Chang, J., Title, P.O., Cowman, P.F., Sallan, L., Friedman, M., Kaschner, K., Garilao, C., Near, T.J., Coll, M. & Alfaro, M.E. (2018). An inverse latitudinal gradient in speciation rate for marine fishes. Nature, 559, 392–395.

Rabosky, D.L., Donnellan, S.C., Grundler, M. & Lovette, I.J. (2014). Analysis and Visualization of Complex Macroevolutionary Dynamics: An Example from Australian Scincid Lizards. Systematic Biology, 63, 610–627.

Rabosky, D.L., Grundler, M., Anderson, C., Title, P.O., Shi, J.J., Brown, J.W., Huang, H. & Larson, J.G. (2014). BAMMtools: an R package for the analysis of evolutionary dynamics on phylogenetic trees. Methods in Ecology and Evolution, 5, 701–707.

Rabosky, D.L., Mitchell, J.S. & Chang, J. (2017). Is BAMM Flawed? Theoretical and Practical Concerns in the Analysis of Multi-Rate Diversification Models. Systematic Biology, 66, 477–498.

Rabosky, D.L., Title, P.O. & Huang, H. (2015). Minimal effects of latitude on present-day speciation rates in New World birds. Proceedings of the Royal Society of London. Series B: Biological Sciences, 282, 20142889.

Redding, D.W. & Mooers, A.O. (2006). Incorporating Evolutionary Measures into Conservation Prioritization. Conservation Biology, 20, 1670–1678.

Ricklefs, R.E. (2006). Global variation in the diversification rate of passerine birds. Ecology, 87, 2468–2478.

Rohde, K. (1992). Latitudinal Gradients in Species Diversity: The Search for the Primary Cause. Oikos, 65, 514–527.

Rosindell, J., Cornell, S.J., Hubbell, S.P. & Etienne, R.S. (2010). Protracted speciation revitalizes the neutral theory of biodiversity. Ecology Letters, 13, 716–727.

Sanderson, M.J. & Donoghue, M.J. (1996). Reconstructing shifts in diversification rates on phylogenetic trees. Trends in Ecology and Evolution, 11, 15–20.

Schluter, D. (2016). Speciation, Ecological Opportunity, and Latitude. The American Naturalist, 187, 1–18.

Silvestro, D., Schnitzler, J. & Zizka, G. (2011). A Bayesian framework to estimate diversification rates and their variation through time and space. BMC Evolutionary Biology, 11, 311.

Stadler, T. (2011). Simulating Trees with a Fixed Number of Extant Species. Systematic Biology, 60, 676–684.

Steel, M. & Mooers, A. (2010). The expected length of pendant and interior edges of a Yule tree. Applied Mathematics Letters, 23, 1315–1319.

Zink, R.M., Klicka, J. & Barber, B. (2004). The tempo of avian diversification during the Quaternary. Philosophical Transactions of the Royal Society B: Biological Sciences, 359, 215–220.

